# Polyphenol rewiring of the microbiome reduces methane emissions

**DOI:** 10.1101/2024.10.22.619724

**Authors:** Bridget B. McGivern, Jared B. Ellenbogen, David W. Hoyt, John A. Bouranis, Brooke Stemple, Rebecca A. Daly, Samantha H. Bosman, Matthew B. Sullivan, Ann E. Hagerman, Jeffrey P. Chanton, Malak M. Tfaily, Kelly C. Wrighton

## Abstract

Methane mitigation is regarded as a critical strategy to combat the scale of global warming. Currently, about 40% of methane emissions originate from microbial sources, which is causing strategies to suppress methanogens—either through direct toxic effects or by diverting their substrates and energy—to gain traction. Problematically, current microbial methane mitigation knowledge derives from rumen studies and lacks detailed microbiome-centered insights, limiting translation across ecosystems. Here we utilize genome-resolved metatranscriptomes and metabolomes to assess the impact of a proposed methane inhibitor, catechin, on greenhouse gas emissions for high-methane-emitting peatlands. In microcosms, catechin drastically reduced methane emissions by 72-84% compared to controls. Longitudinal sampling allowed for reconstruction of a novel catechin degradation pathway involving Actinomycetota and *Clostridium*, which break down catechin into smaller phenolic compounds within the first 21 days, followed by degradation of phenolic compounds by *Pseudomonas_E* from days 21 to 35. These genomes also co-expressed hydrogen-uptake genes, suggesting that hydrogenases may act as a hydrogen sink during catechin degradation, depriving methanogens of substrates. This was supported by decreased gene expression in hydrogenotrophic and hydrogen-dependent methylotrophic methanogens under catechin treatment. We also saw reduced gene expression from genomes inferred to be functioning syntrophically with hydrogen-utilizing methanogens. We propose that catechin metabolic redirection effectively starves hydrogen-utilizing methanogens, offering a potent avenue for curbing methane emissions across diverse environments including ruminants, landfills, and constructed or managed wetlands.

## Introduction

Reducing greenhouse gas concentrations in the atmosphere is necessary to limit the mean global temperature increase to 2°C above pre-industrial levels. Methane (CH_4_) is a greenhouse gas with nearly 30-times the warming potential of carbon dioxide (CO_2_) [1]. There is increasing recognition of the urgent need to limit CH4 emissions to reach climate goals [2], with initiatives like the Global Methane Pact seeking to reduce methane emissions by 20% of 2020 levels by 2030 [3]. Current estimates suggest 60% of CH4 is anthropogenic, while 40% is derived from natural sources [4]. Wetlands represent the largest natural source of methane emissions [4,5]. In these ecosystems, methane (CH_4_) is produced by methanogenic archaea through three pathways, utilizing hydrogen and CO_2_ (hydrogenotrophic), acetate (acetoclastic), or methylated compounds (methylotrophic) [6]. The substrates for these methanogenic pathways are largely produced by other members of the soil microbiome through organic matter decomposition, highlighting the complex microbial interactions underlying methane production in wetland ecosystems.

Ruminant systems have proven tractable for testing and identifying methane inhibition strategies [7], yet translation of these approaches to wetland ecosystems remains challenging. Methane mitigation efforts in ruminants have been either direct, specifically targeting methanogens, or indirect, rerouting methanogenic substrates [8]. For instance, the direct inhibition of the critical methanogenesis enzyme methyl-coenzyme reductase by 3-nitrooxypropanol has been shown to reduce ruminant methane emissions by 30% [9]. Indirect strategies involve shifting reducing equivalents to alternative hydrogen sinks instead of methanogenesis [10]. For example, adding mixtures of plant polyphenol to the animal diet has been shown to variably reduce enteric methane [11]. In particular, the polyphenolic flavonoid catechin (**Fig. 1A**) reduced *in vitro* rumen methane emissions by 20%, potentially serving as an alternative hydrogen sink scavenging electrons from methanogens [12]. However, the microbial pathways and metabolic networks underlying this inhibition remain unexplored, limiting our understanding of the mechanism and its potential application to other ecosystems.

**Figure 1.**
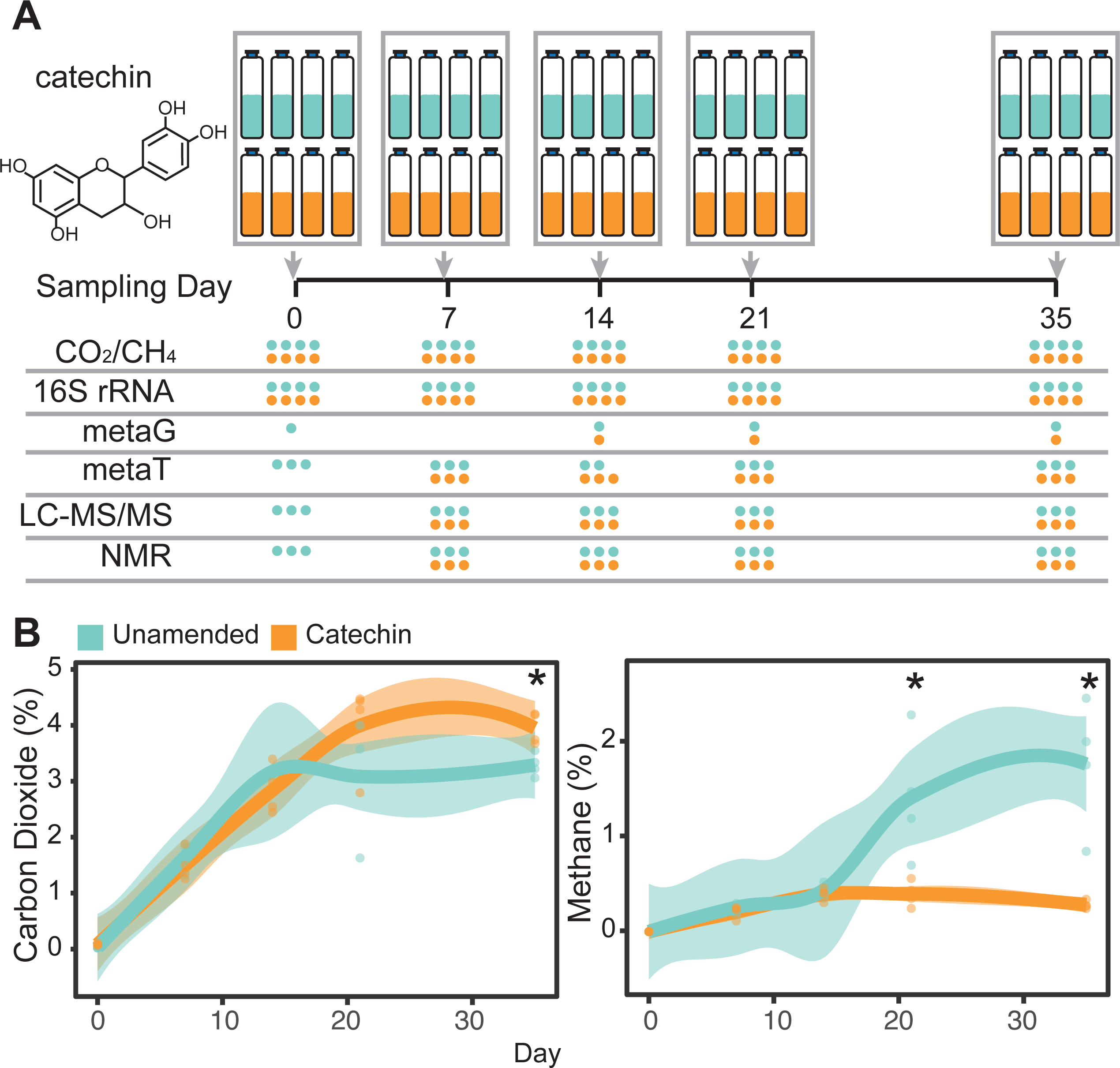
Peat microcosm experiment designed to investigate the impact of catechin amendment on microbial carbon cycling. **(A)** Unamended (teal) and catechin-amended (orange) peat microcosms were constructed and destructively sampled over 35 days at the indicated time points. Headspace carbon dioxide (CO_2_) and methane (CH_4_) were sampled, DNA was extracted for 16S rRNA gene amplicon sequencing and metagenomes (metaG), RNA was extracted for metatranscriptomes (metaT), and metabolites were extracted and analyzed for metabolomes (LC-MS/MS, NMR). Circles correspond to the number of replicates for each data type in the final dataset, colored by treatment. **(B**) Headspace carbon dioxide (left) and methane (right) concentrations over time in the microcosms. Concentrations is given as percent (%) volume. Timepoints with significant differences by treatment are noted with an asterisk (*, Kruskal-Wallis test, p<0.05). Smoothed curves represent the average gas concentration (n=4), with individual replicates plotted as points, and the shaded area represents the 95% confidence interval.

The translation of methane mitigation strategies across ecosystems is further complicated by variations in microbiome structure and function. For example, ruminant systems exhibit lower diversity in methanogen taxonomy and metabolisms than what is observed in wetland soils [13]. For example, acetoclastic methanogenesis is an important pathway in terrestrial systems [6,14], but relatively insubstantial in rumen microbiomes [13,15]. Furthermore, the rumen host depends on its microbiome to generate short chain fatty acids for its energy, constraining the degree to which rumen microbiome metabolism can be rewired [16]. In contrast, wetland microbiomes do not seem to have the same metabolic constraints, potentially allowing for more flexible microbial metabolic rewiring in response to interventions.

Here, we used peat fen microcosms to interrogate the impact of catechin amendment on microbial carbon cycling and greenhouse gas emissions. Building on our previous work that tracked microbial players and enzymes involved in catechin degradation in anoxic wetland soil [17], we now link these metabolisms to carbon greenhouse gas emissions. Our research goals in this current work were (1) to measure the impact of catechin amendment on CO_2_ and CH_4_ production in the microcosms, and (2) to resolve the microbiome metabolic response to catechin amendment. Combining amplicon, genome-resolved metatranscriptome, metabolite, and emission data, we reconstructed how catechin amendment remodeled the peat microbiome carbon cycle. These results provide a mechanistic view of a hydrogen sink in a terrestrial system and highlight additional control points for methane inhibition that could be targeted by future inhibition strategies.

## Materials and Methods

### Peat Sampling

Peat cores were taken from a wetland fen in Stordalen Mire (Abisko, Sweden; 68° 22ʹ N, 19° 03ʹ E) in July 2016 using an 11-cm-diameter push corer. Cores were sectioned in the field, and the depth section corresponding to 9-19 cm below surface was used in this study. The sectioned core was immediately placed into a plastic bag and sealed. Cores were transported on ice from the field to the research station and stored at −20 °C until shipment on dry ice, then returned to −20 °C storage until microcosm construction.

### Incubation Set-Up

The 40 microcosms were sampled destructively over time at 5 timepoints (**Fig. 1**). To construct the fen incubation microcosms, the frozen fen core was thawed at room temperature for 1 hour. Roughly 6g of thawed peat was added to sterilized glass Balch tubes (**Supplementary Data 1**). The tubes were then sealed with a sterile butyl stopper and aluminum crimp. Tubes were flushed with N_2_ gas for 5 minutes, then 10 mL of sterile anoxic media was added. The media consisted of (per liter): 0.25 g ammonium chloride, 0.60 g sodium phosphate, 0.10 g potassium chloride in sterile water with N_2_ headspace. Then, unamended microcosms received an additional 1mL of the sterile anoxic media, while catechin-amended microcosms received 1mL of a sterile, anoxic 15mg/mL (+)-catechin hydrate (CAS# 7295-85-4, Sigma-Aldrich) aqueous solution with N_2_ headspace, resulting in a final catechin concentration of 4.4mM. After the media and amendments were added, the tubes were vortexed to create a slurry and flushed with N_2_ gas for 10 more minutes. At this time, the day 0 microcosms were harvested and the remaining tubes were placed in a dark incubator at 19°C, representing field temperature [18].

Microcosms were harvested at days 0, 7, 14, 21, and 35. To destructively sample the microcosms, headspace gas was first taken from the tubes (described below, *Gas Measurements*). Then, tubes were uncapped, and the slurry was decanted into a 15mL falcon tube and centrifuged for 10min at 16,000xg. From the clarified supernatant, two 1mL aliquots were stored at -80°C in 1.7mL microcentrifuge tubes for metabolite-based NMR and LC-MS analysis. Another 1mL was taken and used immediately to measure pH using an Accumet AB150 Benchtop pH Meter (Fisher, **Supplementary Data 1**). The remaining liquid was discarded, and the pellet was immediately stored at -80°C for nucleic acid extraction for DNA and RNA.

### Gas Measurements

To measure headspace CO_2_ and CH_4_, 10 mL headspace was removed from microcosms using a gas tight Hamilton syringe and stored in glass 6.9 mL Exetainer vials. Vials were stored at room temperature until the end of the experiment and shipped at ambient temperature to Florida State University for GC-FID analysis. CO_2_ and CH_4_ were measured on a Shimadzu 8A gas chromatograph with a carbosphere packed column operated at 140°C. CO_2_ was converted to CH_4_ for Flame ionization analysis (FID) by running it across a methanizer. Samples were quantified relative to calibrated air gas standards.

### Nucleic Acids Extraction

DNA and RNA was extracted from the frozen pellets using the ZymoBIOMICS DNA/RNA Miniprep Kit. For the lysis step, the pellet was resuspended in 750 µL kit lysis solution, transferred to the provided lysis bead tubes, and lysed with a FastPrep-24 at 5 meters/second for 20 seconds. Extracted DNA and RNA was quantified using the Qubit HS dsDNA and HS RNA kits, respectively. Extracted DNA was stored at -20°C until sequencing, and extracted RNA was stored at -80°C until sequencing.

### 16S rRNA Gene Sequencing and Analysis

16S rRNA gene amplicon libraries were prepared from the extracted DNA using barcoded Earth Microbiome Project (EMP) primers 515F [19] (GTG**Y**CAGCMGCCGCGGTAA) and 806R [20] (GGACTAC**N**VGGGTWTCTAAT) and following the EMP amplification protocol [21]. Libraries were normalized using SequalPrep Normalization plate kits (Invitrogen), and pooled. Libraries were sequenced on an Illumina MiSeq at the Center for Microbial Exploration Sequencing Center at University of Colorado-Boulder.

A range of 60,542 to 243,678 16S rRNA gene amplicon read pairs were obtained per sample (**Supplementary Data 2**). Reads were demultiplexed and analyzed within QIIME2 (2021.2) [22] using DADA2 [23] to produce a table of amplicon sequence variants (ASVs) by sample. Taxonomy was assigned using a naïve Bayes sklearn classifier trained with the GTDB-Tk species representative genomes (release 207) [24]. Feature tables were randomly subsampled to a total of 50,000 read counts per sample (**Supplementary Data 2**).

A Bray-Curtis dissimilarity matrix was calculated from ASV abundances using vegan (v2.6-4) [25]. Differences by treatment and time were assessed using the adonis2 function in vegan (v2.6-4) [25].

### Metagenome Sequencing, Assembly, and Binning

Metagenomes were obtained from the seven samples indicated in **Fig. 1**, in addition to five other enrichment samples constructed from unamended Stordalen Mire bog peat microcosms (n=2) and Stordalen Mire fen peat amended with condensed tannin (n=3). These additional five samples were used purely for metagenome sequencing and genome recovery. Metagenome libraries were prepared using the Tecan Ovation Ultralow V2 DNA-Seq kit and sequenced on a NovaSeq6000 system (Illumina, v.1.5 chemistry, S4 flow cell, 2 × 150 bp) at the Genomics Shared Resource Facility at the University of Colorado Anschutz Medical Campus.

Fastq files were trimmed with sickle (v1.33) [26]. We used several assembly and binning strategies to recover metagenome-assembled genomes (MAGs). This workflow is depicted in **Supplementary Fig. 1.** Briefly,

(1) Individual assemblies. Each metagenome sample was assembled individually with MEGAHIT (v1.2.9) [27] using the following flags: --k-min 31 --k-max 121 --k-step 10. Coverage information was determined for contigs >2,500 base pairs using bbmap [28] and a BAM file was generated using samtools (v1.9) [29]. These contigs were binned using metaBAT2 (v1.2.9) [30]. MAG quality was assessed using checkM2 (v0.1.3) [31]. MAGs with completion >50% and contamination <10% were retained as medium and high quality (MQ/HQ) MAGs [32].
(2) Some individual samples were iteratively assembled. Trimmed metagenome reads from individual samples were mapped against MQ/HQ MAGs generated from the individual assemblies and reads that did not map were assembled as described above. Contigs from these assemblies were binned as described above. This was repeated until no/very few MQ/HQ MAGs were recovered.
(3) We combined trimmed reads from different samples and coassembled them with MEGAHIT using the flags mentioned above. Contigs from these assemblies were binned using the combined reads to generate coverage. For two coassemblies, multi-sample coverage was used in metaBAT2, using the coverage of the contigs from the individual reads that went into the coassembly.

The MQ/HQ MAGs recovered from these efforts were dereplicated at 99% nucleotide identity using dRep (v2.6.2) [33]. The cluster winners were dereplicated at 99% nucleotide identity with 13,290 MAGs from Stordalen Mire [34]. MAG taxonomy was inferred using the Genome Taxonomy Database Toolkit (GTDB-tk v2.3.0 r214) [35]. MAGs were annotated using DRAM (v1.4.4) [36] and CAMPER (v1) [34].

### Metatranscriptome Sequencing and Analysis

Metatranscriptome libraries were prepared and sequenced at the Joint Genome Institute. Plate-based RNA sample prep was performed on the PerkinElmer Sciclone NGS robotic liquid handling system using the FastSelect 5S/16S/23S for bacterial rRNA depletion kit (Qiagen) with RNA blocking oligo technology to block and remove rRNA from 100ng of total RNA input. An Illumina sequencing library was then created from the fragmented and rRNA-depleted RNA using the TruSeq Stranded Total RNA HT sample prep kit (Illumina) following the protocol and with 10 cycles of PCR for library amplification. The prepared libraries were quantified using KAPA Biosystems’ next-generation sequencing library qPCR kit and run on a Roche LightCycler 480 real-time PCR instrument. Sequencing of the flow cell was performed on the NovaSeq sequencer (Illumina) using NovaSeq XP V1.5 reagent kits, S4 flow cell, following a 2x151 indexed run recipe.

Raw metatranscriptome reads were quality trimmed and adapters removed using bbduk [28] with the following flags: k = 23 mink=11 hdist=1 qtrim=rl trimq=20 minlength=75. Each sample was randomly subsampled to 50,000,000 pairs of trimmed reads, and the reads were mapped against the database of 99% dereplicated MAGs using bowtie2 (v2.4.5) [37] with the following flags: -D 10 -R 2 -N 1 -L 22 -i S,0,2.50. The output SAM file was converted to BAM using samtools and filtered using the reformat.sh script in the bbtools package using: idfilter=0.97 pairedonly=t primaryonly=t. Mapped reads were counted using htseq-count (v.0.13.5) [38] with the following flags: -a 0 -t CDS -i ID --stranded=reverse. Read counts were filtered to remove counts <5 and were converted to geTMM [39] in R. We determined the median number of genes with non-zero geTMM per MAG to be 22, and thus filtered out genes from MAGs with less than 22 genes “on” in a sample. Scripts for processing metatranscriptome count data are provided in GitHub (see **Data Availability**).

Beta-diversity in the metatranscriptome was assessed three ways: first, a Bray-Curtis dissimilarity matrix was calculated from gene-level geTMM abundances using vegan (v2.6-4) [25]; second, gene-level geTMM abundances were summed at the MAG level per sample, and a Bray-Curtis dissimilarity matrix was calculated from this MAG table using vegan (v2.6-4) [25]; third, gene-level geTMM abundances were summed at the annotation level (using DRAM annotations from ko_id, cazy_best_hit, and camper_id columns) per sample, and a Bray-Curtis dissimilarity matrix was calculated from this annotation table using vegan (v2.6-4) [25]. Differences by treatment and time were assessed using the adonis2 function in vegan (v2.6-4) [25].

### Classifying MAG gene expression responses to catechin

We developed a classification scheme for the MAGs based on their gene expression across the treatments at each timepoint. This was developed using two metrics: (1) the percentage of genes expressed in both unamended and catechin-amended microcosms (%both) and (2) the difference in the percentage of genes expressed only in unamended and only in catechin amended microcosms (Δunique) (**Supplementary Fig. 2**). Using these two values and defined thresholds assigned based on their distribution across the MAGs that recruited transcripts (**Supplementary Fig. 2**), MAGs were assigned to one of 6 categories at each timepoint: (1) resistant, %both >= 26, |Δunique|=<25; (2) responsive, %both < 26, |Δunique|=<25; (3) sensitive, %both < 26, Δunique >=25; (4) stimulated, %both < 26, Δunique =< -25; (5) lost function, %both >= 26, Δunique >=25; (6) gain function, %both >= 26, Δunique =< -25. Additional details are provided in **Supplementary Table 1**.

### Metabolism Curation

(1) *Methanogen metabolism curation.* To curate methanogen metabolism, DRAM-annotated MAGs were screened following the method published in [14]. In brief, MAGs were first confirmed as methanogens via the presence of genes encoding the Mcr and Hdr complexes, with consideration of taxonomy and other methanogenic genes. MAGs were then investigated for substrate use potential and classified as either (1) hydrogenotrophic (encoding genes for the Wood-Ljungdahl pathway, Mtr complex subunits, and relevant hydrogenases) (2) acetoclastic (member of the *Methanosarcinia* encoding the ACS/CODH complex and either acetyl-CoA synthetase or acetate kinase/phosphate acetyltransferase) or (3) methylotrophic (encoding genes for a three-component methyltransferase system including at least one substrate:corrinoid methyltransferase, termed *mtxB*). MAGs were further screened for recently characterized methylotrophic genes missing from DRAM using a curated BLASTP approach. MAGs seen to encode multiple pathways were denoted as multifunctional. Metranscriptomic data was then queried for these MAGs expression of methanogenic genes. These genes were categorically grouped into “pathways” based on their belonging to (1) the Wood-Ljungdahl pathway, (2) a methylotrophic methyltransferase system, (3) their function in acetate utilization or (4) their belonging to a methanogenic protein complex (e.g. Mcr complex). MAGs were classified as actively utilizing distinct substrates in reactors at each time point by the following rules:

a. hydrogenotrophic if actively expressing at least one gene belonging to the Wood-Ljungdahl pathway and at least one relevant hydrogenase gene
b. methylotrophic if actively expressing a substrate-specific *mtxB* gene (substrate:corrinoid methyltransferase)
c. acetoclastic if found to be expressing the aforementioned acetate genes (*acs* or *ack/pta*).

Active *Methanotrichales* MAGs were considered obligate acetoclasts, otherwise multi-functional methanogens were noted to utilize multiple substrates if multiple rules were met by a single MAG. MAGs found to express no methanogenesis genes at a timepoint, or genes insufficient to meet any of these rules (e.g. active but not classifiable to a pathway), were ignored for metatranscriptomic analyses focused on pathway expression.
(2) *Hydrogenase curation.* To curate potential and expression of genes predicted to encode hydrogenases, we searched the MAG database with Hidden Markov Models (HMMs) developed for [NiFe], [FeFe], and [Fe] hydrogenases [40], using the provided scores as cut-offs. We pulled the amino acid sequences of passing genes and placed them in a phylogenetic tree with HydDB [41] representatives to assign genes to hydrogenase groups (**Supplementary Fig. 3**). Hydrogenase group was determined using the placement of genes of interest relative to the HydDB representatives. For genes where the placement was not clear, we manually submitted the sequence to the HydDB webserver. We did not refine the [FeFe] Group A sequences into subgroups. We inferred hydrogenase directionality using HydDB “Activity” for each hydrogenase group, where “H_2_-evolving” hydrogenases produce H_2,_ “H_2_-uptake (unidirectional)” hydrogenases consume H_2_, “Bidirectional” indicates both H_2_ production and consumption, and “Electron-bifurcation” could indicate both H_2_ production and consumption.
(3) *Carbon cycle curation.* To assign MAGs to roles in the carbon cycle, we used combinations of MAG taxonomy and DRAM annotations. See **Supplementary Data 3** for the rules used to call pathways.

### LC MS/MS

The frozen 1mL aliquot of microcosm supernatant was thawed at 4 °C. Following thawing, the sample was centrifuged to remove any particles that may have formed during the freeze-thaw process. Next, each sample was split into two vials (1 ml each), one for hydrophilic interaction liquid chromatography (HILIC) and the other for reverse-phase (RP) liquid chromatography as described previously [42,43]. These samples were completely dried down using a Vacufuge plus (Eppendorf, USA). Following this, the samples were resuspended in a solution of 50% Acetonitrile and 50% water for HILIC and a solution of 80% water and 20% HPLC grade methanol for RP.

For the chromatography step, a Thermo Scientific Vanquish Duo ultra-high performance liquid chromatography system (UHPLC) was employed at the University of

Arizona Analytical & Biological Mass Spectrometry Facility. Extracts were separated using a Waters ACQUITY HSS T3 C18 column for RP separation and a Waters ACQUITY BEH amide column for HILIC separation. Samples were injected in a 1 μL volume on column and eluted as follows: for RP the gradient went from 99% mobile phase A (0.1% formic acid in H2O) to 95% mobile phase B (0.1% formic acid in methanol) over 16 minutes. For HILIC the gradient went from 99% mobile phase A (0.1% formic acid, 10 mM ammonium acetate, 90% acetonitrile, 10% H2O) to 95% mobile phase B (0.1% formic acid, 10 mM ammonium acetate, 50% acetonitrile, 50% H2O). Both columns were run at 45 °C with a flowrate of 300 μL/min.

Spectral data collection was performed at the University of Arizona Analytical & Biological Mass Spectrometry Facility using a Thermo Scientific Orbitrap Exploris 480. The instrument operated with a spray voltage of 3500 V for positive mode (for RP) and 2500 V for negative mode (for HILIC) using the H-ESI source. Both the ion transfer tube and vaporizer temperature were both 350 °C. Compounds were fragmented using data-dependent MS/MS with HCD collision energies of 20, 40, and 80.

Data analysis was conducted using Compound Discoverer 3.3.2.31 software (Thermo Fisher Scientific) following the untargeted metabolomics workflow. Briefly, the spectra were first aligned followed by a peak picking step. Putative elemental compositions of unknown compounds were predicted using the exact mass, isotopic pattern, fine isotopic pattern, and MS/MS data using the built in HighChem Fragmentation Library of reference fragmentation mechanisms. Metabolite annotation was performed using and in-house database built using 1200 reference standards, spectral libraries and compound databases. First, fragmentation scans, retention time and ion mass of unknown compounds were compared with those in the in-house database. Second, fragmentation scans (MS2) searches in mzCloud were performed, which is a curated database of MSn spectra containing more than 9 million spectra and 20000 compounds. Third, predicted compositions were obtained based on mass error, matched isotopes, missing number of matched fragments, spectral similarity score (calculated by matching theoretical and measured isotope pattern), matched intensity percentage of the theoretical pattern, the relevant portion of MS, and the MS/MS scan. The mass tolerance used for estimating predicted composition was 5 ppm. Compounds level of annotation was assigned according to the Metabolomics Standards Initiative [44] as follows: Level 1: compounds with exact match to a standard reference compound in our in-house library, Level 2: compounds with full match to online spectral databases using mzCloud database (based on MS2 spectra matching) and Level 3: for compounds with only a molecular formula (generated through the Predicted Compositions node).

To reduce noise and collinearity and remove artifactual peaks from the dataset, a custom in house script was used to identify in source fragments and ions resulting from neutral mass loss. Briefly, ions were first binned based on their retention time using a retention time tolerance window of 0.005 seconds. Next, compounds which co-eluted were examined for correlations in their raw intensity and only ions which had a linear correlation across all samples greater than 0.98 were retained. Next, the MS2 spectra of the heaviest ion (the parent ion) was searched for each candidate fragment ion with mass similarity of less than 5 ppm. Only candidate fragments ions present in the MS2 of the parent ion were retained. Lastly, the resulting candidate list was manually inspected through MS2 matching, MS2 annotation (based on neutral mass losses), and by examining the correlation patterns to determine the validity of the candidate fragment ions. Validated fragment ions were removed from the data set. Additionally, compounds with predicted compositions containing halogen ions were removed from the dataset as it is unlikely these compounds would exist naturally in peat soils and are thus likely artefacts of analysis.

LC MS/MS metabolite peak areas were used to assess metabolome beta-diversity. A Bray-Curtis dissimilarity matrix was calculated from peak areas of level 1 and level 2 HILIC and RP metabolites using vegan (v2.6-4) [25]. Differences by treatment and time were assessed using the adonis2 function in vegan (v2.6-4) [25].

### NMR

The frozen 1mL aliquot of microcosm supernatant was thawed on ice. Once thawed, 180 µL supernatant was combined with 2,2-dimethyl-2-silapentane-5-sulfonate- d_6_ (DSS-d_6_) in D_2_O (20 µL, 5 mM) and thoroughly mixed prior to transfer to 3 mm NMR tubes. NMR spectra were acquired on a Bruker Neo spectrometer operating at 18.8T (1H ν0 of 800.30 MHz) equipped with a 5mm Bruker TCI/CP HCN (inverse) cryoprobe with Z-gradient at a regulated temperature of 298.0 K. The 90° ^1^H pulse was calibrated prior to the measurement of each sample. The one-dimensional ^1^H spectra were acquired using a nuclear Overhauser effect spectroscopy (noesypr1d) pulse sequence with a spectral width of 20.1 ppm and 2048 transients. The NOESY mixing time was 100ms and the acquisition time was 4s followed by a relaxation delay of 1.5 s during which presaturation of the water signal was applied. The 1D ^1^H spectra were manually processed, assigned metabolite identifications and quantified using Chenomx NMR Suite 9.0. Time domain free induction decays (65536 total points) were zero filled to 131072 total points prior to Fourier transform, followed by exponential multiplication (0.3 Hz line-broadening), and semi-automatic multipoint smooth segments baseline correction. Chemical shifts were referenced to the ^1^H methyl signal in DSS-d6 at 0 ppm. Metabolite identification was based on matching the chemical shift, J-coupling and intensity of experimental signals to compound signals in the Chenomx, HMDB and custom in-house databases (**Supplementary Data 5**). Quantification was based on fitted metabolite signals relative to the internal standard (DSS-d_6_). Signal to noise ratios (S/N) were checked using MestReNova 14.3 with the limit of quantification equal to a S/N of 10 and the limit of detection equal to a S/N of 3.

### Statistics

All data analyses and visualization were done in R (v.4.3.2) [45] with the following packages: stats, ggplot2 (v.3.4.4) [46], tidyr (v.1.3.0) [47], dplyr (v.1.1.4) [48], pheatmap (v.1.0.12) [49], and vegan (v2.6-4) [25].

## Results

### Catechin amendment alters methane production

We constructed anoxic microcosms with a peat slurry using material from a fen in Stordalen Mire, an artic peatland in Sweden. Over 35 days, the unamended and 4.4 mM (+)-catechin amended peat microcosms were sampled at key timepoints (**Fig. 1A**) for headspace CO_2_ and CH_4_, DNA for 16S amplicon sequencing and metagenomes, RNA for metatranscriptomes, and metabolites for LC MS/MS and NMR analyses. The resultant gas measurements and multi-omics longitudinal dataset were then used to assess how catechin altered greenhouse gas production, as well as microbial responses to catechin amendment.

We first assessed the impact of catechin amendment on CO_2_ and CH_4_ production. Based on *in vitro* bovine rumen fluid experiments [12], we expected to see catechin amendment cause a reduction in net methane production. Our data confirmed this hypothesis, illustrating net methane production was significantly reduced relative to the unamended control by 72% and 84% after 21 and 35 days, respectively (**Fig. 1B, Supplementary Data 1**). In contrast, net CO_2_ production was not reduced with catechin amendment, and instead was significantly higher by 20% after 35 days (**Fig. 1B, Supplementary Data 1**). Together, the CO_2_ and CH_4_ data suggest that microbial carbon decomposition occurred in the presence of catechin, thereby producing CO_2_, yet the downstream products appeared not to feed methanogens. Presumably, the decreased CH_4_ is not due to an overall toxicity to the microbial community, but instead from a revision of the metabolic network away from methane emission pathways and towards different compounds. To test this, we next investigated the underlying microbiome dynamics both in organismal relative abundance and gene expression over the time series.

### Microbial community gene expression changes with catechin amendment

We tracked the microbial communities using genome-resolved metatranscriptomes. This effort leveraged a database of microbial metagenome-assembled genomes (MAGs) compiled from two sources. First, we recovered MAGs from 623 Gbp of metagenomic sequencing derived from the microcosms. Using a combination of assembly and binning approaches, we generated 815 medium and high-quality MAGs that were dereplicated into 300 MAGs at 99% nucleotide identity (**Supplementary Fig. 1**, **Supplementary Data 2**). Second, we used a database of 13,290 MAGs recovered from 8-years of field-derived metagenomes more broadly sampled at the source field site (Stordalen Mire) [34]. We dereplicated the 300 microcosm MAGs with this field MAG set at 99% nucleotide identity, leading to a final database of 2,302 dereplicated, medium and high-quality MAGs (**Supplementary Fig. 4, Supplementary Data 2**). Of the microcosm MAGs, 56 were dereplication cluster losers, 15 were dereplication cluster winners, and 229 were singletons (**Supplementary Fig. 4, Supplementary Data 2**). This illustrates the depth of microbial diversity in terrestrial systems and the value of enrichment experiments to recover under sampled members of field microbial communities.

From the final set of 2,302 MAGs, 982 MAGs recruited transcripts in the metatranscriptome (metaT, **Supplementary Data 3**). These MAGs contributed a total of 1,242,169 genes, spanning 8,340 functional IDs (see *Methods*). To capture these levels of data resolution, we analyzed beta-diversity in the metatranscriptome three ways: (1) at a gene level (metaT gene), (2) at a genome-level by summing gene expression across a MAG (metaT MAG), and (3) at a function level by summing genes at the level of their annotation (metaT annotation; ex. KO id, dbCAN id). These analyses revealed time and catechin amendment were significant drivers of beta-diversity in the three metatranscriptome measures, 16S rRNA gene amplicon community, and the LC MS/MS metabolome (**Supplementary Fig. 5**).

To deduce which of these microbial measurements was most responsive to polyphenol amendment, we compared the Bray-Curtis distances from the catechin-amended samples to the unamended control samples at each time point (**Fig. 2A**). This revealed that metabolite and gene-level metatranscriptome data had the highest Bray-Curtis distances between unamended and catechin-amended samples at each timepoint, on average 1.6- and 2-fold higher than at the 16S rRNA gene, metaT MAG, and metaT annotation levels (**Fig. 2A**). Notably, the metaT gene level had higher Bray-Curtis distances than the metaT MAG level and metaT annotation level. Overall, while the identity (metaT MAG level) and functionality (annotation level) of the active microbial community changed with catechin amendment, gene expression changes within MAGs were the most dynamic. We interpret these findings to indicate a degree of functional redundancy in the microbial community.

**Figure 2.**
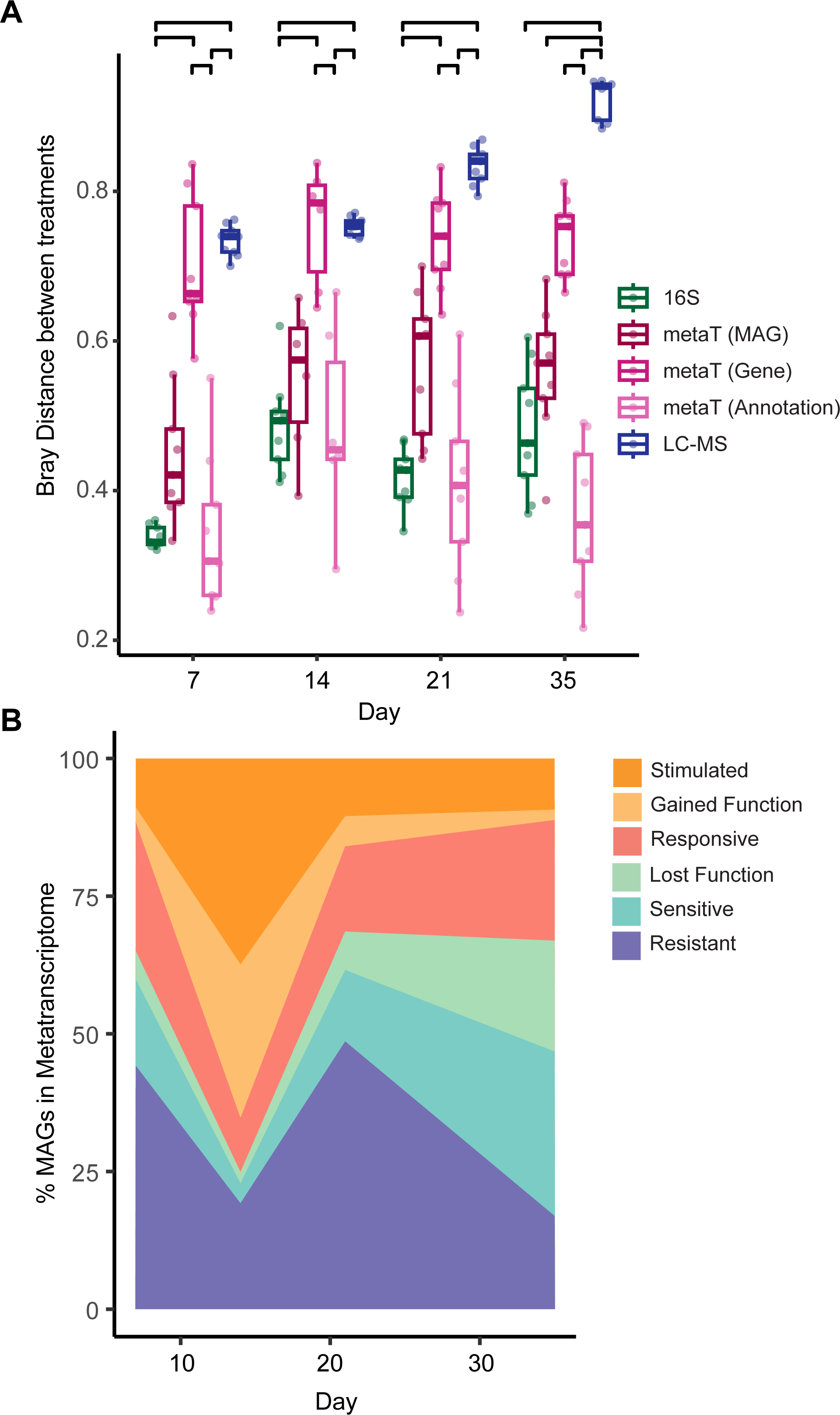
Dynamics of microbiome datasets. **(A)** Bray-Curtis distances were calculated at each timepoint between catechin-amended and unamended replicates for 16S rRNA amplicon (16S) samples, metatranscriptome (metaT), and metabolite (LCMS) samples. MetaT samples were analyzed at three levels: metaT abundance of individual genes (Gene), metaT abundance of genes summed at the MAG level (MAG), and metaT abundance of gene functions, summing genes at the annotation level (Annotation). Significant differences are noted at the top with brackets (Kruskal-Wallis with post-hoc Dunn’s, Benjamini-Hochberg adjusted p<0.05). The lower and upper boxplot edges represent the 25th and 75th percentiles, respectively, and the middle line is the median. The whiskers extend from the median by 1.5X the interquartile range. (**B**) The proportion of MAGs transcriptionally expressing genes in the six response categories at each timepoint. See **Supplementary Fig. 4** for the data at each time point, and *Methods* for classification methods.

Next, we wanted to identify which MAGs altered their gene expression with catechin amendment, and how they changed. MAGs were assigned to one of six categories (see *Methods,* **Supplementary Fig. 2**). The first category, *resistant*, was assigned when the genes expressed by a MAG were essentially unaffected by catechin amendment. Conversely, the second category, *responsive*, was assigned when MAGs expressed different genes in each treatment. Third and fourth, MAGs were assigned as *stimulated* or *sensitive* if most of their genes were expressed exclusively in catechin or unamended microcosms, respectively. Finally, the last two categories *gained function* and *lost function* were assigned when MAGs expressed a set of genes in both treatments in addition to a unique set of genes in catechin or unamended microcosms, respectively.

We looked at the percentage of MAGs that were assigned to each category of all MAGs that recruited transcripts from day 7 to day 35. By day 14, 65% of active MAGs were in the *stimulated* or *gained function* category, while just 5% were in the *sensitive* or *lost function* categories (**Fig. 2B**). From day 21 to day 35, the proportion of MAGs classified as *sensitive* or *lost function* increased from 20% to 50%, while MAGs classified as *stimulated* or *gained function* decreased to less than 16% of the community. This latter trend mirrors the CH_4_ data, where net CH_4_ production is impacted after 21 days. Thus, we next wanted to examine the microbial processes and players in these distinct phases.

### Reconstruction of catechin degradation pathway

Both the LC MS/MS and NMR metabolomics showed steady decreases in catechin abundance and concentration, respectively, to baseline by day 21, suggesting this was the period of catechin biodegradation (**Fig. 3A, Supplementary Data 5**). To understand this catechin removal, we next sought to reconstruct the catechin degradation pathway using metabolites and gene expression. Catechin degradation begins with the conversion of catechin into taxifolin [50,51], though the responsible enzyme remains unknown. Then, taxifolin degradation proceeds through two known routes characterized in microbial isolates (**Fig. 3A**, Route 1-2). In one route [52], the enzyme flavanone- and flavanonol- cleaving reductase (FCR) cleaves the C-ring of taxifolin, generating a dihydrochalcone product which is cleaved by the enzyme phloretin hydrolase (PHY). The products of this reaction are 3,4-dihydroxyphenyllactic acid and phloroglucinol. In the other route [53], taxifolin undergoes ring-contraction via the enzyme chalcone isomerase (CHI) to form the auronol alphitonin. Alphitonin is then hypothesized to be degraded into 3,4- dihydroxyphenylacetic acid and phloroglucinol [54].

**Figure 3.**
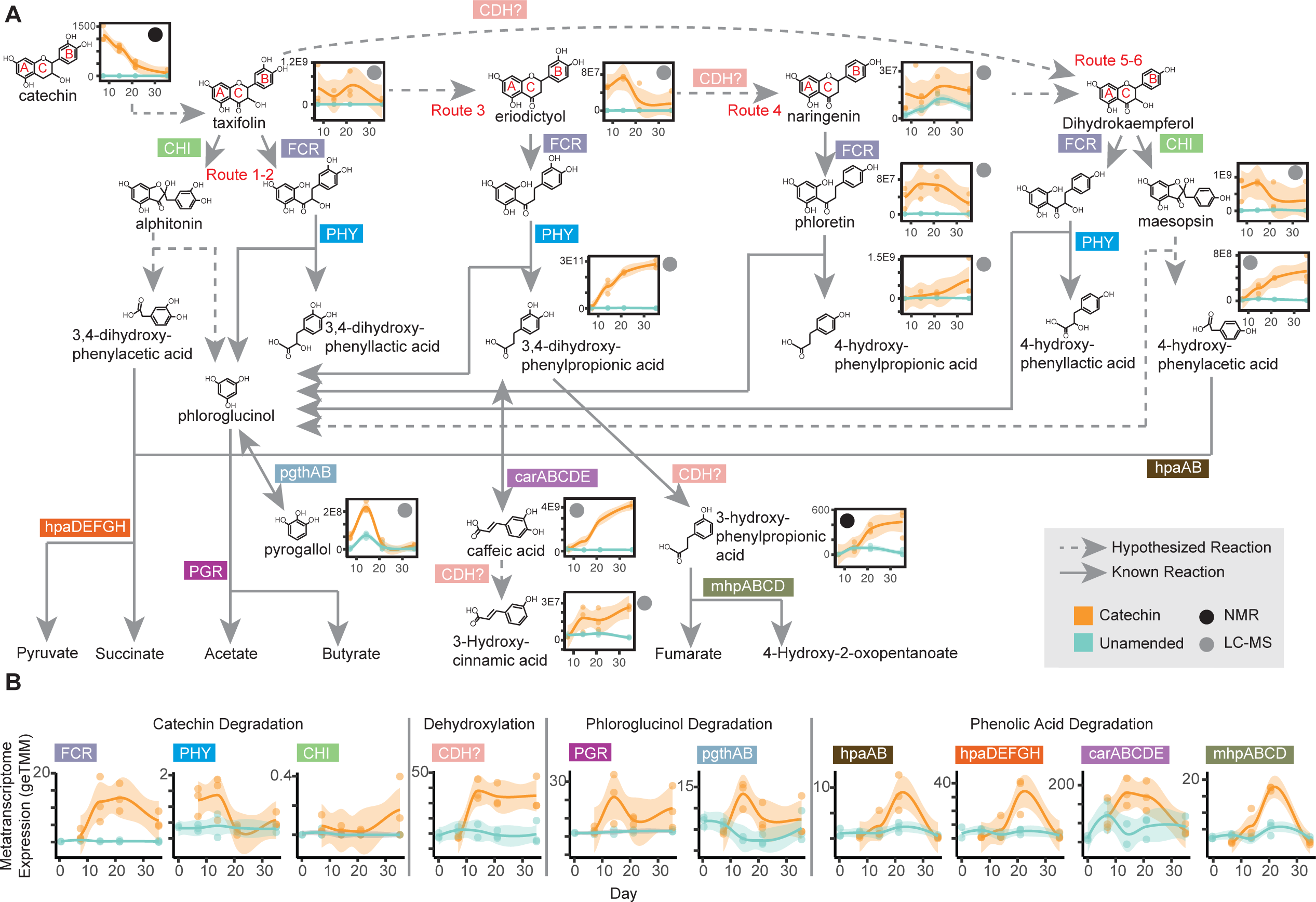
Catechin degradation pathway reconstructed from metabolite and metatranscriptome data. (**A**) Reconstruction of catechin and phenolic acid degradation pathways using metatranscriptome and metabolite data. Flavonoid rings are labelled A, B, and C to correspond to the text. Hypothesized reactions are noted with dotted arrows, while known reactions are shown with solid arrows. Metatranscriptome-detected genes encoding enzymes for the reactions are noted on the arrows. Metabolite dynamics of LC-MS or NMR detected metabolites are shown adjacent to compound name, from days 7-35, with detection method noted with a grey and black dot, respectively. Concentrations of NMR metabolites are given in µM, while normalized peak area is given for LC-MS metabolites. (**B**) Gene expression profiles of catechin and phenolic acid degrading genes across metatranscriptomes. Orange bars label enzymes by the part of catechin degradation with which they are involved. Timepoints (days 0- 35) are given on the x-axis, and the summed gene expression (geTMM) is given on the y-axis. In **A** and **B,** smoothed curves represent the average metabolite abundance and metatranscriptome expression, respectively (n=3), with individual replicates plotted as points, and the shaded area represents the 95% confidence interval.

Interestingly, taxifolin was the only intermediate from the known catechin degradation pathway detected in our metabolite data, present exclusively in catechin-amended microcosms through day 21 (**Fig. 3A, Supplementary Data 5**). However, we detected metabolites from three other potential routes of degradation, suggesting a broader metabolic fate for catechin than previously realized. One potential pathway appears to proceed from dehydroxylation at C-3 of the C-ring of taxifolin, producing the flavanone eriodictyol, which is detected at day 7 and day 14 in catechin-amended microcosms (**Fig. 3A**, Route 3). Then, successive action of FCR and PHY can lead to phloroglucinol and 3,4-dihydroxyphenylpropionic acid; the latter increases in the metabolite data from day 14 on.

The second potential route could proceed from eriodictyol by dehydroxylation of C-3 of the B-ring to the flavanone naringenin. Naringenin can be converted to phloretin by FCR, and phloretin can be converted to 4-hydroxyphenylpropioic acid and phloretin by PHY (**Fig. 3A**, Route 4). Naringenin, phloretin, and 4-hydroxyphenylpropionic acid are all detected in our metabolomes. Finally, the third route could proceed from dehydroxylation of taxifolin at C-3 of the B-ring to generate the flavanonol dihydrokaempferol (**Fig. 3A**, Route 5-6). FCR and PHY can act on dihydrokaempferol [52], though we do not detect the intermediate products of these enzymes. Instead, we detected the auronol maesopsin, which could result from CHI-mediated ring-contraction of dihydrokaempferol. Maesopsin was hypothesized to degrade into phloroglucinol and the metabolome- detected compound 4-hydroxyphenylacetic acid.

Given these potentially novel catechin degradation pathways inferred from metabolomics data, we next sought to evaluate whether transcript data supported this metabolite-informed degradation scheme. Mirroring what was expected from metabolite data, expression of genes for FCR and PHY peaked at days 14 and 21 (**Fig 3B, Supplementary Data 6**). We detected low expression of CHI, which fits with sparse detection of CHI products in the metabolite data. We observed high expression of genes annotated as catechol dehydroxylases (CDH). A clade of these enzymes has been shown to dehydroxylate catechin at the C-4 position of the A-ring [55]. Therefore, we postulate these may be involved in the conversion of eriodictyol to naringenin and/or taxifolin to dihydrokaempferol via unexplored biochemistry.

Beyond catechin degradation, we detected gene expression and metabolite support for degradation of the downstream phenolic products from catechin. At day 14, gene expression for two pathways converting key intermediate phloroglucinol peaked (**Fig 3B, Supplementary Data 6**). The first pathway, initiated by the enzyme phloroglucinol reductase (PGR), degrades phloroglucinol to acetate and butyrate [56]. In the second, phloroglucinol is reversibly converted to pyrogallol by the enzyme pyrogallol transhydroxylase (pgthAB) [57]. In support of this gene expression, pyrogallol peaks in our metabolite data at day 14 (**Fig. 3A, Supplementary Data 5**). Following day 21, we saw increased expression of genes encoding enzymes in pathways for degradation of 3,4-dihydroxyphenylacetic acid (hpaDEFGH), 3,4-dihydroxyphenylpropionic acid (carABCDE), and 4-hydroxyphenylacetic acid (hpaAB, **Fig 3B, Supplementary Data 6**). We hypothesize CDH could also act on these phenolic products, generating dehydroxylated products. For example, 3,4-dihydroxyphenylpropionic acid could be dehydroxylated to the detected metabolite 3-hydroxyphenylpropionic acid (**Fig. 3A, Supplementary Data 5**). We additionally detected gene expression of 3- hydroxyphenylpropionic acid degradation (mhpABCD, **Fig 3B, Supplementary Data 6**). Collectively, metabolite and metatranscriptome data illustrate that catechin degradation occurs through day 21 via newly proposed routes, followed by degradation of the phenolic products from day 21 to 35.

### Catechin degradation by novel lineages

Having identified the catechin and phenolic acid degradation pathways most likely functioning in our microcosms, we next wanted to identify the microorganisms involved. In total, we found FCR, PHY, and/or CHI transcripts corresponding to 13 MAGs across three phyla, leading us to infer they were the active catechin degraders (**Fig. 4, Supplementary Data 6**). These MAGs belonged to nine genera (*Mycobacteria*, *Clostridium, Clostridium_I*, *Pelorhabdus*, undescribed Actinobacterial genera CAQPS01, Chersky-299, and RBG-16-64-13, undescribed Bacillota_A genus JAGFXR01, and a novel genus in the Bacillota_C family CTSoil-080), with most gene expression coming from JAGFXR01 (FCR genes) or *Clostridium* (PHY and CHI genes, **Supplementary Fig. 6**). In contrast to just a few taxa dominating catechin degradation gene expression, genes for phloroglucinol transformation (PGR, pgthAB) were more broadly expressed across the microbial community. In fact, 57 MAGs from 13 phyla, including nearly all catechin- degrading MAGs expressed genes for phloroglucinol degradation (**Fig. 4, Supplementary Data 6**). In addition to the undescribed lineages expressing genes for catechin degradation, a further 20 MAGs involved in phloroglucinol degradation were from alphanumeric genera. The majority of phloroglucinol degradation gene expression came from *Clostridium* and JAGFXR01 (**Supplementary Fig. 6**), further marking these lineages as critical players in catechin-degradation.

**Figure 4.**
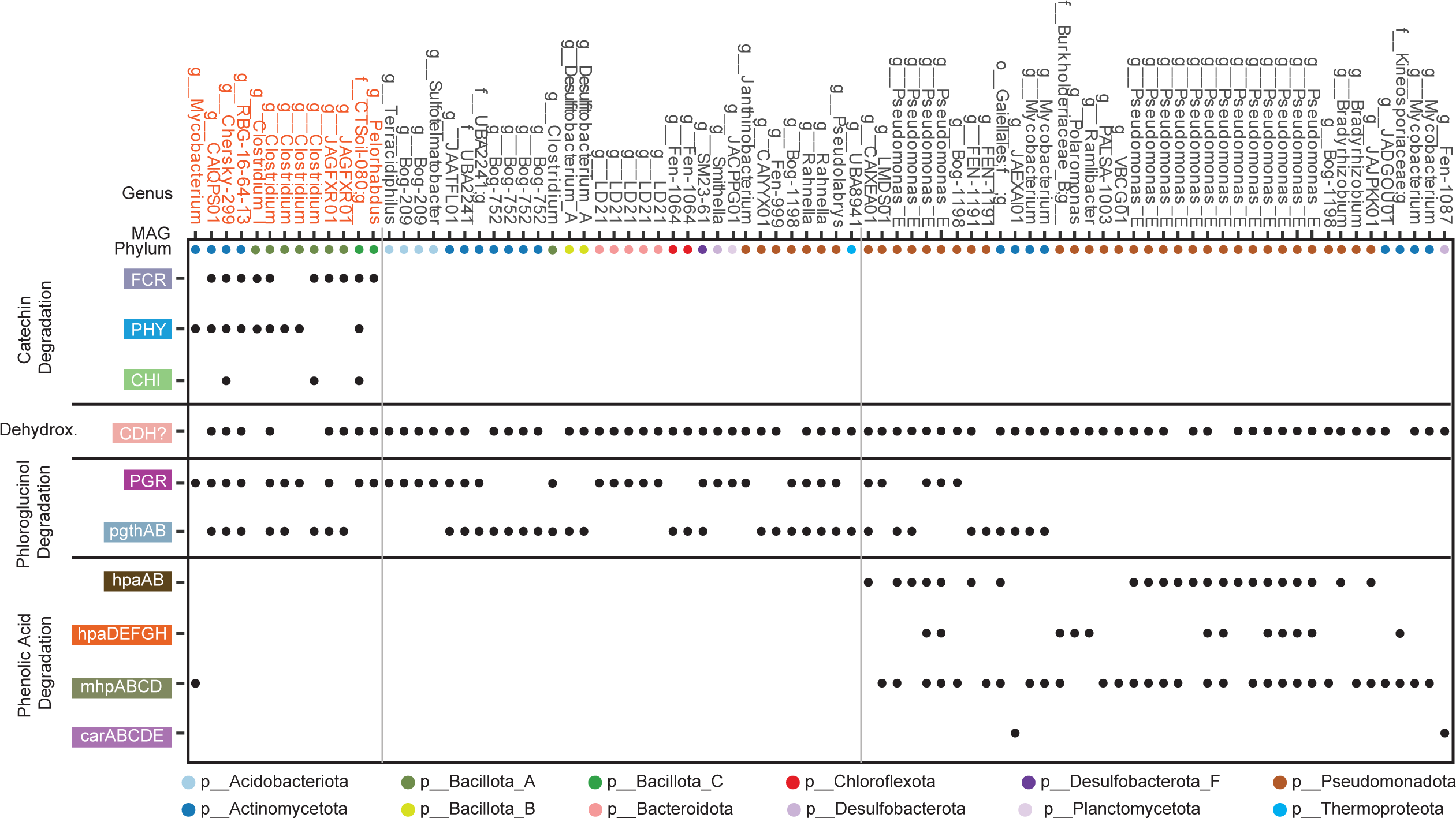
Diverse microbial lineages involved in catechin and phenolic acid degradation. The microbial MAGs expressing genes encoding enzymes for catechin and phenolic acid degradation in the catechin-amended reactors. MAG genus is listed at top, and colored dots correspond to MAG phylum. Grey vertical lines delineate MAGs broadly involved in catechin degradation (highlighted in red text), dehydroxylation (dehydrox.), phloroglucinol degradation, and phenolic acid degradation. A black dot denotes a MAG expressed the given gene. Only MAGs expressing genes for at least two enzymes are shown; for full data, see **Supplementary Data 6.** For pathways with multiple genes, at least 50% of genes needed to be expressed to be considered positive.

The degradation of catechin-derived phenolic acids could be traced to 41 MAGs in 18 genera. This included 17 MAGs in the genus *Pseudomonas_E (***Fig. 4, Supplementary Fig. 6, Supplementary Data 6)**. Interestingly, no catechin-degrading MAGs expressed genes for degradation of the phenolic acid products, suggesting these are distinct degrading functional groups. CDH gene expression was detected from 73 MAGs from 42 genera, further underscoring the need to validate the role of these cryptic enzymes which were widely expressed (**Fig. 4, Supplementary Data 6)**. In all, we showed the main players in catechin and phloroglucinol degradation were MAGs in *Clostridium* and JAGFXR01, followed by degradation of phenolic acids by *Pseudomonas_E*.

### Methanogen gene expression was impacted by catechin addition

Next, we investigated metabolic processes at work following catechin and phenolic acid degradation, where most transcriptionally active genomes were classified as *lost function* or *sensitive* with catechin amendment. Given the dramatic impact of catechin addition on methane production, we first focused on methanogen activity. Gene expression was detected from 50 methanogen MAGs spanning 9 genera across the experimental conditions and timepoints (**Fig. 5A**). The relative number of metatranscriptome reads recruited to methanogen MAGs was not significantly different between catechin-amended and unamended microcosms at any time point, although recruitment trended lower in catechin-amended (**Supplementary Fig. 7**). We observed peak gene expression from most methanogen MAGs at days 21 and 35 in the unamended microcosms, but not in the catechin-amended microcosms (**Fig. 5A**). This trend mirrored CH_4_ production data (**Fig. 1B**). Further, 96% of active methanogen MAGs were classified as *lost function* or *sensitive* at day 35 (n=46, **Fig. 5A**), reinforcing the negative impact of catechin amendment on methanogen activity.

**Figure 5.**
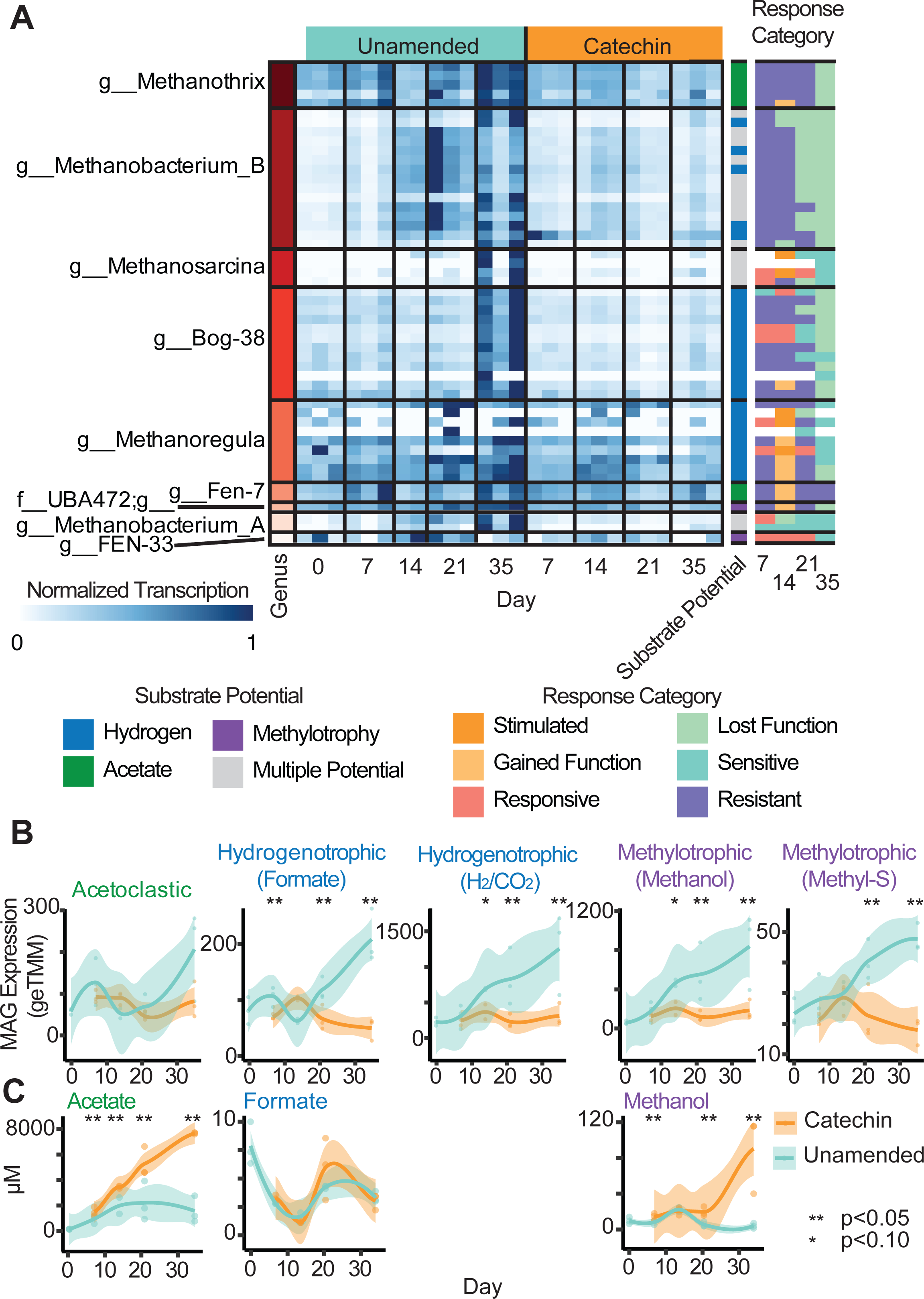
Methanogen metatranscriptome activity decreased in catechin-amended microcosms. **(A)** Normalized metatranscriptome expression of methanogen MAGs (rows) in each metatranscriptome sample (column). Dark blue corresponds to the highest expression (normalized transcription =1) detected in a sample for each MAG, with subsequent samples colored accordingly. Methanogenic substrate potential is colored at right (**Supplementary Data 7)**, and methanogen MAG response classification is given across the time series at far right. In MAG response classification, white corresponds to MAGs without detectable metatranscriptome expression. **(B)** Total methanogen MAG expression grouped by methanogenic pathway (**Supplementary Data 7)**. (**C**) Concentrations of methanogenic substrates detected by NMR in µM. In B-C, smoothed curves represent the average metatranscriptome expression and concentration, respectively(n=3), with individual replicates plotted as points, and the shaded area represents the 95% confidence interval. Timepoints with significant differences between unamended and catechin-amended are marked with asterisks (Kruskal-Wallis test, * p-value <0.10, ** p-value <0.05).

The methanogen MAGs had metabolic potential for methanogenesis via acetoclastic, hydrogenotrophic, and methylotrophic pathways (**Fig. 5A**). To identify whether a particular methanogenesis pathway was impacted by catechin amendment, we analyzed methanogen MAG-level transcription profiles (**Supplementary Fig. 7, Supplementary Data 7**). The main acetoclastic gene expressing lineages were MAGs in *Methanosarcina* and Fen-7 (family *Methanotrichaceae*). Methylotrophic gene expression came from MAGs in the undescribed family UBA472 (Methanomassiliicoccales) and *Methanobacterium_B*. The UBA472 MAG expressed genes for methyl-sulfur compound utilization (*mtsDFH*), while *Methanobacterium_B* MAGs expressed genes for methanol (*mtaBCA*). MAGs from *Methanobacterium_A*, *Methanobacterium_B*, *Methanoregula* and Bog-38 (*Ca.* Methanoflorens [58,59]) expressed genes for hydrogenotrophic methanogenesis. Some Bog-38 and *Methanoregula* MAGs expressed formate dehydrogenase, suggesting they were using formate, while other *Methanoregula*, Bog- 38, *Methanobacterium_B*, and *Methanobacterium_A* MAGs appeared to be using H_2_ and CO_2_. Six *Methanobacterium_B* MAGs co-expressed genes for methylotrophic and hydrogenotrophic methanogenesis. These genera aligned with the known active methanogens in Stordalen Mire, suggesting the key methane cycling members recapitulated field communities in our microcosms [14].

At a pathway level, acetoclastic methanogen MAG expression was not significantly impacted by catechin amendment (**Fig. 5B**). This may be because acetate is a product of phloroglucinol degradation (**Fig. 4A**) and thus acetoclastic methanogens were not substrate limited. This is evidenced by the metabolite data, where acetate concentrations were nearly 4-times higher in the catechin-amended microcosms than in the unamended at day 35 (**Fig. 5C**). As methane production and gene expression by acetoclastic methanogens did not spike in response to acetate build up, there may be other constraints on methanogen physiology in these microcosms, for example trace element availability [60].

Conversely, both formate and H_2_/CO_2_ pathways for hydrogenotrophic methanogen gene expression were significantly lower in catechin-amended metatranscriptomes (**Fig. 5B**). Formate concentrations were not significantly different between treatments over time (**Fig. 5C**), indicating other microorganisms outcompeted the methanogens for the formate in the catechin-amended microcosms. Methylotrophic methanogenesis gene expression was also significantly reduced in the catechin-amended microcosms (**Fig. 5B**). Specifically, we saw reduction of methanol utilization (*mtaBCA*) by MAGs in the *Methanobacteriales* and several pathways for diverse methylated compounds by MAGs in the *Methanomassiliicoccales* under catechin amendment (**Supplementary Fig. 7, Supplementary Data 7**). NMR metabolites revealed an increase in methanol over time in catechin-amended microcosms, but not in the unamended (**Fig. 5C**). Unlike acetate, methanol is not an expected degradation product of catechin. Methanol can be produced from pectin demethylation [61] or acetone degradation [62], but gene expression was not significantly different between treatments for these genes (**Supplementary Fig. 8)**. Therefore, we postulate decreased methanogen consumption led to methanol accumulation. Furthermore, the *Methanomassiliicoccales* and *Methanobacterium_B* are thought to be obligate and facultative hydrogen-dependent methylotrophs, respectively [63–65]. Taken together, the decrease in hydrogenotrophic and hydrogen-dependent methylotrophic methanogenesis indicates decreased hydrogen levels may be the source of reduced methane production in the catechin reactors.

### Hydrogen metabolism increased in catechin amended microcosms

Given hydrogen consuming methanogens were most impacted in our microcosms, we next investigated hydrogenase gene expression across treatments and time points. Employing hydrogenase hidden Markov models (HMMs) and phylogenetic trees, we identified 3,480 genes predicted to encode hydrogenase catalytic subunits in our MAG database (**Supplementary Data 8**). Of these genes, 1,000 were expressed in the metatranscriptome spanning 28 hydrogenase subgroups (**Supplementary Fig. 3, Supplementary Data 8**).

Hydrogenases can be classified by their directionality: H_2_ consuming, H_2_ producing, and both H_2_ consuming and producing (bidirectional, bifurcating) [66]. Using these classifications, we observed significantly higher gene expression for H_2_ uptake hydrogenases in catechin amended microcosms at days 14, 21, and 35 (**Fig. 6A**), concurrent with when CH_4_ concentrations were lower. Similarly, gene expression for bidirectional hydrogenases was higher at days 14 and 21 in catechin amended relative to unamended microcosms (**Fig. 6A**). Gene expression for bifurcating hydrogenases from non-methanogens was significantly higher at day 35 in unamended microcosms (**Fig. 6A**). Therefore, we confirmed an increase in hydrogen consumption in catechin amended microcosms, supporting the idea that the catechin degradation pathway may be a hydrogen sink.

**Figure 6.**
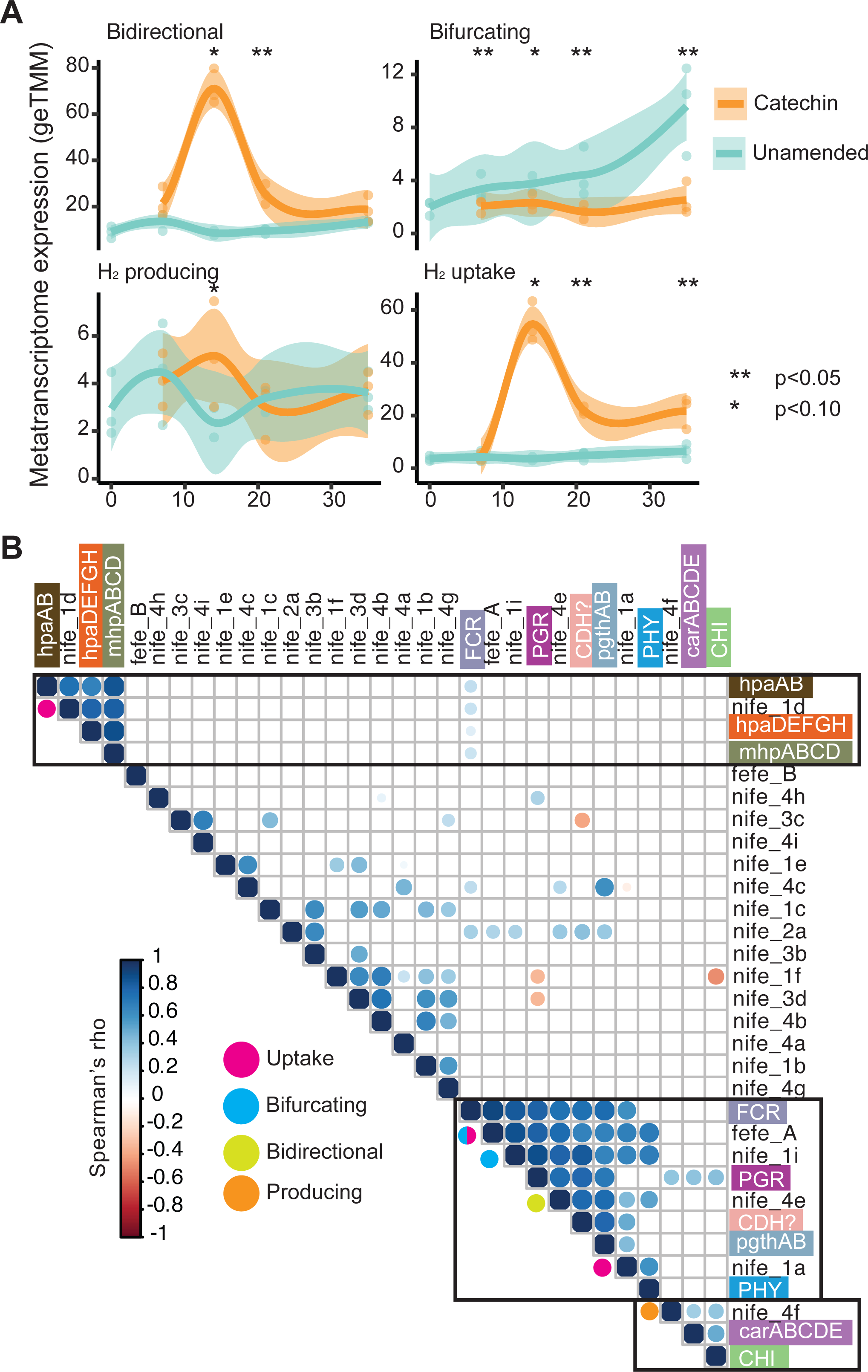
Increased hydrogen metabolism in catechin amended microcosms. **(A)** Gene expression of hydrogenases from non-methanogen MAGs grouped by their hydrogen use. Smoothed curves represent the average metatranscriptome expression (n=3), with individual replicates plotted as points, and the shaded area represents the 95% confidence interval. Timepoints with significant differences between unamended and catechin-amended are marked with asterisks (Kruskal-Wallis test, * p-value <0.10, ** p-value <0.05). (**B**) Correlation matrix showing significant Spearman’s correlations in gene expression between hydrogenase subgroups and catechin/phenolic acid degradation. Significant correlations are marked by a dot, colored by the Spearman’s rho. Boxes are drawn around the correlation clusters with catechin/phenolic degradation genes. Colored dots align on the row of the hydrogenase group, corresponding to hydrogen use.

To link catechin more explicitly to hydrogen metabolism, we correlated expression of catechin/phenolic degradation genes to hydrogenase genes (**Fig. 6B**). There were three distinct correlation clusters. The first cluster included FCR, PHY, PGR, pgthAB, CDH and four hydrogenase subgroups. These hydrogenases include likely H_2_ consuming hydrogenases (some [FeFe]-A, [NiFe]-1a), a bidirectional hydrogenase ([NiFe]-4e), and a bifurcating hydrogenase ([NiFe]-1i). In support of the relationship between these hydrogenases and the genes for catechin and phloroglucinol degradation, most gene expression came from *Clostridium*, JAGFXR01, and JAEXAI01 MAGs (**Supplementary Fig. 6**). The next correlation cluster included gene expression for the phenolic acids produced from catechin and a single hydrogenase group, the [NiFe]-1d hydrogenase. This is a respiratory hydrogenase that consumes H_2_. MAGs from *Pseudomonas_E* contributed most of the gene expression for this hydrogenase (**Supplementary Fig. 6**). The third cluster included CHI and the caffeic acid reductase system (carABCDE), and [NiFe]-4f hydrogenase. This hydrogenase is predicted to produce H_2_ in a respiratory complex using formate as an electron donor; this could be the competing formate-consuming process given methanogenesis from formate was decreased with catechin amendment (**Fig. 5B**). Collectively, we show distinct hydrogenase types were used for H_2_ consumption across catechin and phenolic acid degradation, potentially contributing to the decreased activity of hydrogen-dependent methanogens observed in the late time points in the catechin enrichment.

***Microorganisms across the carbon cycle were negatively impacted by catechin*** Beyond methanogens, we examined the impact of catechin on the broader peat microcosm microbiome. From days 21 to 35, 449 non-methanogen MAGs from 25 phyla were classified as *lost function* or *sensitive* in response to catechin. To better profile the functionalities that were restructured in response to catechin, we inventoried the functions of these MAGs at the genus level from the unamended microcosms. We focused on genera where most (>50%) active MAGs were impacted by catechin, narrowing to 310 MAGs across 65 genera (**Fig. 7 A-B**). The most catechin-impacted genera were Acidobacteriota genus *Terracidiphilus* (37/56 MAGs), Bacteroidota genera LD21 (18/24 MAGs) and *Paludibacter* (18/34 MAGs), and Desulfobacterota genus Fen-1087 (30/35 MAGs) (**Fig. 7A**).

**Figure 7.**
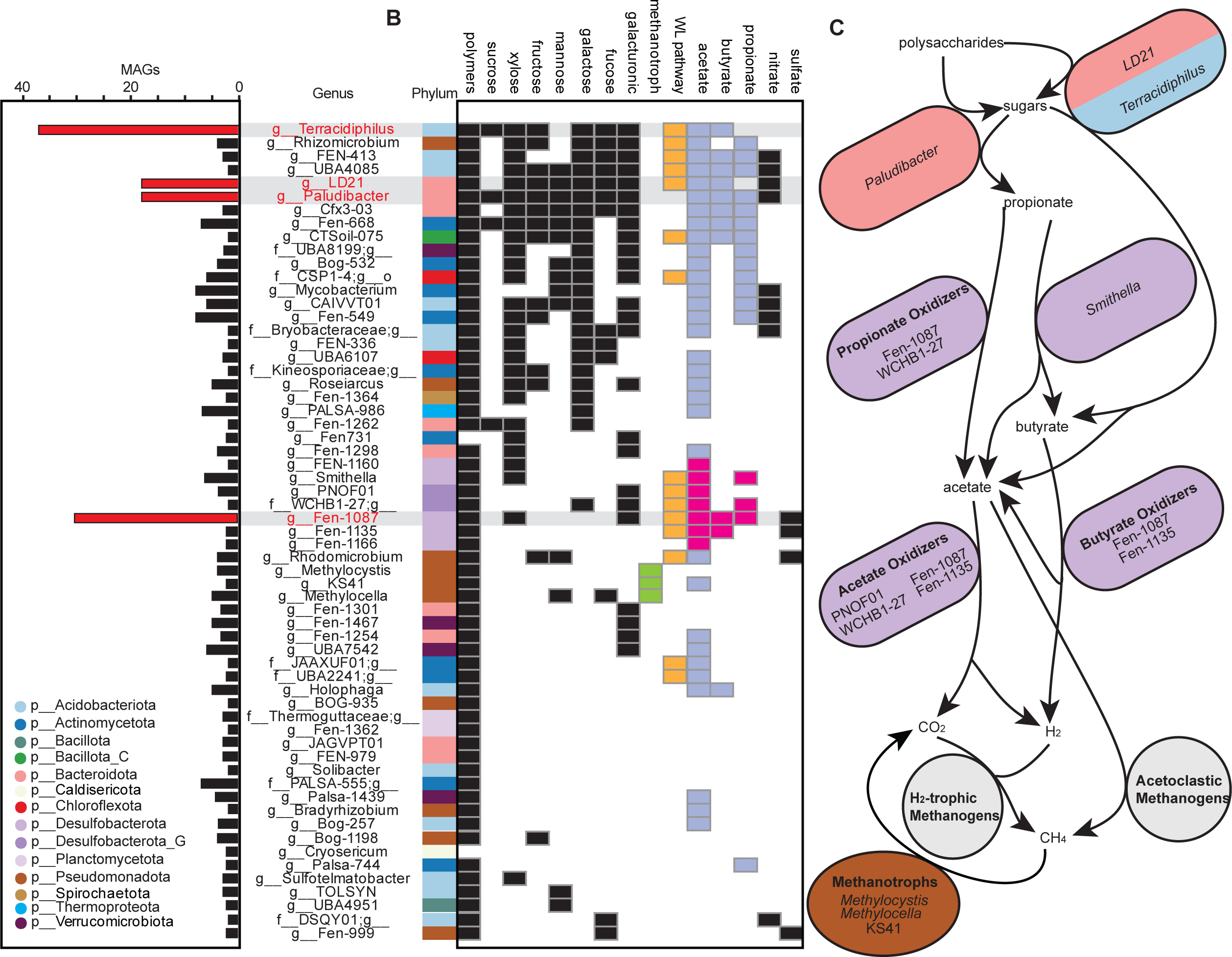
Non-methanogen lineages across the carbon cycle were impacted by catechin amendment. (**A**) The number of MAGs per classified as *lost function* or *sensitive* at days 21 and 35. Only genera with at least two MAG representatives, and greater than 50% active MAGs, were selected for display. Bars highlighted in red correspond to genera referenced in the text. (**B)** Metabolic roles curated from impacted genera gene expression. (**C**) Schematic of carbon cycle in unamended peat microcosms. Cartoon microorganisms are labelled with functional group (bold names) and genus, where able, and colored by phylum.

The impacted genera fit into three groups. Most MAGs, including *Terracidiphilus, Paludibacter,* and LD21, were heterotrophs that expressed genes for polymer degradation and sugar consumption. Carbon was decomposed in these lineages through a mix of fermentation and anaerobic respiration (**Fig. 7A-B**). Second, MAGs from methanotrophic genera *Methylocella, Methylocystis,* and KS41 were negatively impacted, likely because there was significantly less methane available in the catechin-amended microcosms (**Fig. 7B**, green boxes). The third group included MAGs from the phylum Desulfobacterota, including the known syntrophic propionate-oxidizing genus *Smithella* [67] and several genera hypothesized to be propionate (Fen-1087, WCHB1-27), acetate (PNOF01, Fen-1087, WCHB1-27, Fen-1135), and butyrate (Fen-1087, Fen-1135) oxidizing based on gene expression (**Fig. 7B**, pink boxes). In traditional syntrophic relationships, the syntroph oxidizes propionate, acetate, and/or butyrate to H_2_ and CO_2_, where partner hydrogenotrophic methanogens consume these to make the reaction thermodynamically favorable [67] (**Supplementary Table 2**).

Given the breadth of metabolic groups that appeared to be impacted by catechin amendment, we sought to profile the overall differences in expressed carbon cycling genes between the two treatments. Beginning with polysaccharides, we examined expression of carbohydrate-active enzymes (CAZymes). Total CAZyme expression was significantly higher in catechin amended microcosms at day 14, but higher in the unamended at day 35 (**Supplementary Fig. 9**). *Paludibacter* MAGs accounted for an average of 20% of CAZyme expression in the unamended (**Fig. 7C, Supplementary Fig. 9**) but contributed less than 5% of CAZyme expression in the catechin amended. Instead, an average of 34% of CAZyme expression in the catechin-amended microcosms derived from catechin-degrading MAGs in *Clostridium* and *Pseudomonas_E*. Similarly, expression of genes encoding phosphotransferase system enzymes and sugar catabolic pathways was significantly enriched in catechin-amended microcosms at day 14 relative to unamended (**Supplementary Fig. 9**). In support of this, there was significantly less sugars in the catechin-amended microcosms by NMR and LC MS/MS (**Supplementary Fig. 10**). Like for CAZymes, *Paludibacter* MAGs were major sugar gene expressors in the unamended (**Fig. 7C, Supplementary Fig. 9**), but were replaced by *Clostridium* and *Pseudomonas_E* in the catechin-amended microcosms. Together, this data suggests that although overall carbon processing function is maintained, the identity of the active lineages shifted. Importantly, catechin and phenolic acid degrading lineages replaced the dominant carbon degrading MAGs in the unamended control, potentially outcompeting these taxa for substrates.

To understand the impact of this shift in identity of upstream carbon cycling MAGs, we next focused on fermentation pathway gene expression. Likely due to phloroglucinol degradation (**Fig. 4A, Supplementary Fig. 6**), expression of genes for acetate and butyrate fermentation was significantly enriched in the catechin-amended microcosms, with most expression traced to *Clostridium* and *JAGFXR01*. In contrast, propionate fermentation gene expression was significantly higher from days 14 to 35 in the unamended (**Supplementary Fig. 9)**. In the unamended microcosms, 51% of propionate fermentation gene expression was from *Paludibacter* MAGs whereas *Pseudomonas_E* MAGs dominated in the catechin-amended (**Fig. 7C, Supplementary Fig. 8**).

Given the alterations in fermentation pathway gene expression, and that gene expression from putative syntrophic lineages decreased with catechin (**Fig. 7A-B**), we investigated acetate, butyrate, and propionate metabolism gene expression from hypothesized- syntrophic lineage MAGs (**Fig. 7C**, purple cells). Butyrate metabolism gene expression was significantly higher in the unamended microcosms at days 14 and 35 (**Supplementary Fig. 9)**. Gene expression for acetate and propionate metabolism was on average 2-fold and 5-fold higher at day 35, respectively, in the unamended microcosms, though this was not significant (**Supplementary Fig. 9)**. We note that metatranscriptome data alone is not enough to determine if these genes are expressed for oxidation, and that in some cases the genes employed by these lineages are unknown [67].

## Discussion

In this study, we provide the first mechanistic multi-omic insights into how catechin can alter microbiome functionality to inhibit methane emissions in wetland ecosystems. In the paired unamended microbial community, *Paludibacter* and other microorganisms decomposed organic carbon polymers and fermented sugars to propionate. This propionate was likely oxidized to acetate, butyrate, and H_2_ and CO_2_ by purported syntrophic lineages. The H_2_/CO_2_ and acetate were presumably consumed by hydrogenotrophic and acetoclastic methanogens, respectively, and methanol and H_2_ were consumed by methylotrophic methanogens (**Fig. 8**). In the catechin amended microcosms, we propose several new routes for catechin degradation expanding our understanding of polyphenol metabolism in complex microbial communities. These pathways converged at the production of phloroglucinol and several phenolic acids. Phloroglucinol was degraded to acetate and butyrate, and the phenolic acids were degraded into succinate, pyruvate, and fumarate. The organisms that carried out these processes also expressed genes for H_2_ consuming hydrogenases, in addition to CAZymes and sugar degradation genes. We observed the lower trophic levels of the microbiome were inhibited with catechin. Interestingly, acetoclastic methanogens were not significantly impacted by catechin. However, acetate accumulated in the catechin- amended samples without a corresponding increase in acetoclastic methanogenesis, indicating another cryptic process, such as trace element availability, may be limiting these methanogens.

**Figure 8.**
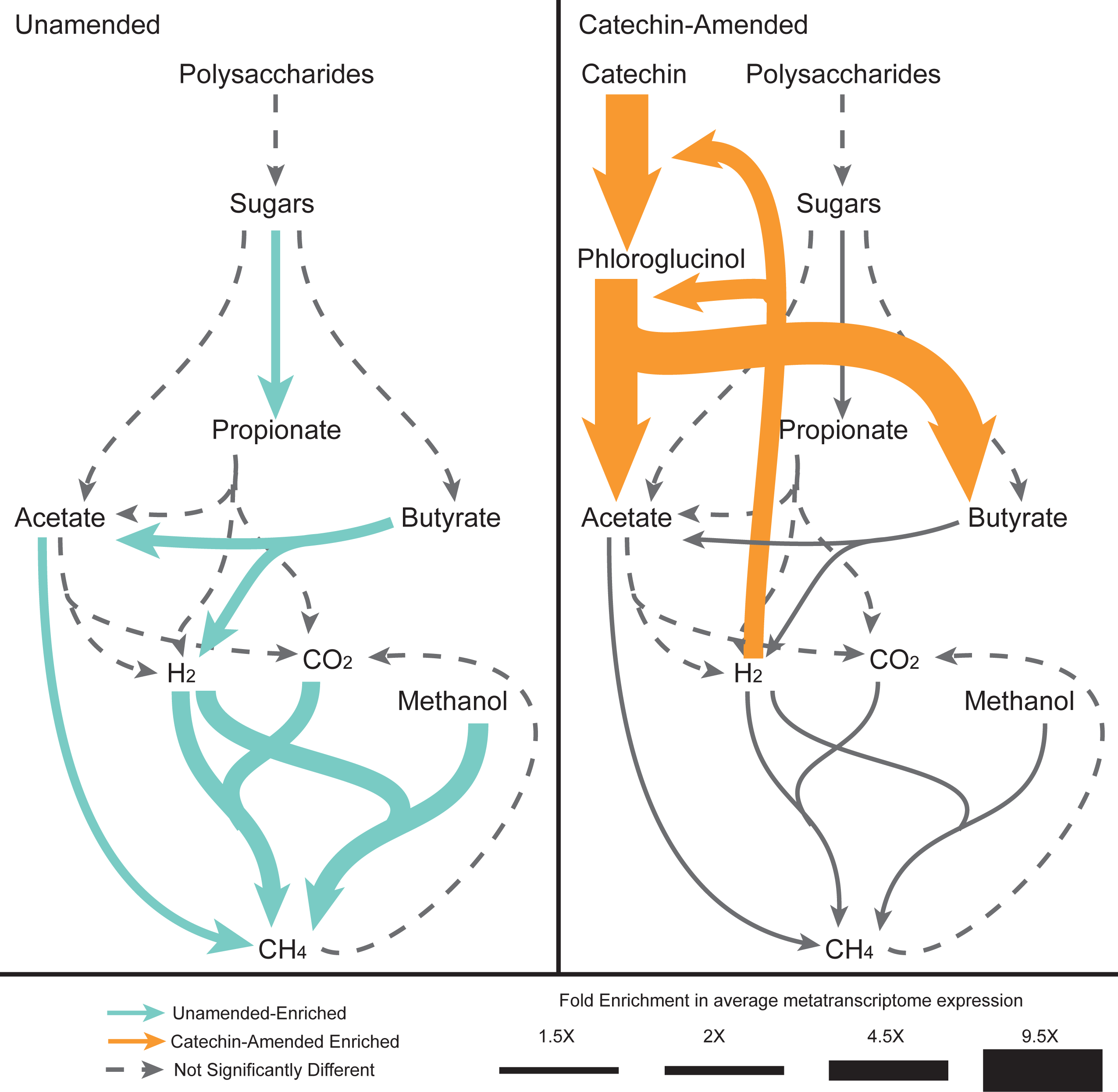
Conceptual model of how catechin could rewire the microbial carbon cycle. Carbon cycle in unamended (left) and catechin-amended (right) peat microcosms. Pathway metatranscriptome expression was assessed with MaAsLin2 (q<0.25), and pathways that were significantly associated with unamended or catechin metatranscriptomes are colored teal and orange, respectively (**Supplementary Data 3**). Arrow thickness corresponds to the average fold enrichment of a pathway. We note a similar rewiring may also occur in the rumen, likely by different taxa, where hydrogen availability modulates methanogenesis.

We postulate catechin inhibits methanogenesis in two ways: (1) as a hydrogen sink, where catechin-degrading microbes increase H_2_ consumption and rob hydrogenotrophic and H_2_-dependent methylotrophic methanogens of hydrogen, and (2) by inhibiting syntrophic relationships, preventing cross-feeding between syntrophic propionate, acetate, and butyrate oxidizing lineages and H_2-_using methanogens. The first scenario may serve as a specific example of ‘biohydrogenation’ of organic matter, a mechanism suggested to limit CH_4_ emissions in *Sphagnum* dominated peatlands [59]. In the second scenario, catechin-degrading microorganisms may outcompete propionate fermenting organisms like *Paludibacter*, reducing propionate pools for propionate oxidizers. It could also result from altered thermodynamics making these oxidation reactions more unfavorable because of changes in substrate and product concentrations following catechin degradation (**Supplementary Table 2**). For example, we observed significant reductions in butyrate metabolism gene expression from putative syntrophic lineages. In this reaction, butyrate is oxidized to acetate, H_2_, and a proton. As acetate concentrations increased and pH decreased with catechin amendment, these conditions likely decreased favorability.

Our findings echo results of catechin amendment in rumen systems. In one study, catechin amendment of *in vitro* rumen fluid reactors led to a proportional decrease in CH_4_ production [12], though the microbial community was not sampled. Our results go a step further to sample how changes in the hydrogen economy can impact the pathway circuitry of methanogenesis *in vitro*. In our study, catechin was over three times more effective at reducing CH_4_ than in the rumen system, limiting CH_4_ emissions by over 70% after 21 days (**Fig. 1B**). This highlights the need to study inhibition strategies across ecosystems. While it is likely not feasible to use catechin amendments in natural systems to shut off methane emissions, here we use this perturbation to highlight control points in the carbon cycle that could inspire targets of other hydrogen scavenging mitigation strategies.

This study represents a first step in linking catechin degradation and methane inhibition. Catechin has been detected in peat extracts from Stordalen Mire [34], however there is little data on catechin concentrations in peat. Therefore, it is unclear if all soils and peatlands will have the capacity for catechin degradation and if there is a minimum concentration of catechin that leads to methane inhibition. Given this was a microcosm study, the communities were separated from live plants and the dynamic inputs of carbon that come from natural systems. Additionally, catechin was only added at the beginning of the experiment, allowing it to be removed by the microbial community. It is unclear if methanogenesis inhibition would continue after the catechin is completely removed, or if the original carbon cycling network would rebound in activity. While our study provides crucial insights, it also highlights areas for future research. The novel lineages we identified as key players in catechin degradation warrant further investigation, potentially leading to the discovery of new enzymatic pathways and metabolic capabilities. Additionally, the complex temporal dynamics we observed suggest the need for longer-term studies to fully understand the ecosystem-level impacts of polyphenol amendments.

## Conclusion

Given the urgent threats of climate change, there is a need to mitigate greenhouse gas release. Several strategies have been identified for ruminants, but they lack mechanistic insight into the cause of the inhibition and the consequence for the entire microbiome. Our findings demonstrate that catechin can significantly inhibit *in vitro* peatland methane emissions, with a more pronounced effect than observed in ruminant systems. While direct catechin amendments may not be feasible for large-scale application in natural systems, our study highlights key control points in the carbon cycle that could inspire targeted, hydrogen-scavenging mitigation strategies. By providing a mechanistic understanding of how catechin alters microbial community function to inhibit methane production, we identified vulnerabilities in the peatland carbon cycle that future strategies could exploit. These insights open new avenues for developing ecosystem-specific approaches to reduce greenhouse gas emissions from high-methane environments. As we continue to face the challenges of climate change, this mechanistic understanding of microbiome-mediated methane inhibition provides a valuable foundation for future research and the development of effective mitigation strategies in diverse ecosystems.

## Acknowledgements

We thank the EMERGE field team for obtaining peat samples and for coordinating sample exchange. We thank Faith Mendoza from the Tfaily lab for her help in preparing the samples for LC MS/MS analysis. We also thank Lawrence Walker and Krishna Parsawar at the University of Arizona Analytical & Biological Mass Spectrometry Facility for their analytical chemistry expertise and data acquisition via LC-MS/ MS.

## Author Contributions- CRediT

**Bridget B. McGivern**: Conceptualization, Data Curation, Formal Analysis, Investigation, Methodology, Project coordination, Visualization, Writing- Original Draft, Writing- review and editing

**Jared B. Ellenbogen**: Data Curation, Writing- review and editing

**David Hoyt**: Data Curation, Investigation, Resources, Writing- review and editing

**John A. Bouranis**: Data Curation, Writing- review and editing

**Brooke Stemple**: Data Curation, Investigation, Writing- review and editing

**Rebecca A. Daly**: Investigation, Project coordination, Writing- review and editing

**Samantha Bosman**: Investigation, Writing- review and editing

**Matthew B. Sullivan**: Funding Acquisition, Writing- review and editing

**Ann E. Hagerman:** Writing- review and editing

**Jeffrey P. Chanton**: Resources, Writing- review and editing

**Malak M. Tfaily**: Funding Acquisition, Investigation, Resources, Writing- review and editing

**Kelly C. Wrighton**: Conceptualization, Funding Acquisition, Project coordination, Resources, Writing- Original Draft, Writing- review and editing

## Data Availability

The 16S rRNA gene amplicon sequencing, metagenomes, metatranscriptomes and metagenome-assembled genomes (MAGs) used in this paper are available at NCBI under BioProjectID PRJNA1137330. See **Supplementary Data 1** (16S rRNA gene sequencing reads), **Supplementary Data 2** (Metagenome reads and MAGs) and **Supplementary Data 3** (metatranscriptome reads) for specific accessions.

MAG Annotations, gene nucleotide, and gene amino acid files are available in Zenodo at (https://doi.org/10.5281/zenodo.13936221). Metatranscriptome mapping files are available in Zenodo at https://doi.org/10.5281/zenodo.13937409. Code for analysis and figure generation can be found on GitHub (https://github.com/WrightonLabCSU/catechin_incubations).

## References

1. Myhre G, Shindell D, Bréon F et al. Anthropogenic and natural radiative forcing. Climate Change 2013: The Physical Science Basis. Contribution of Working Group I to the Fifth Assessment Report of the Intergovernmental Panel on Climate Change. Cambridge University Press, 2013, 659–740.

2. Nzotungicimpaye C-M, MacIsaac AJ, Zickfeld K. Delaying methane mitigation increases the risk of breaching the 2 °C warming limit. Commun Earth Environ 2023;4:1–8.

3. Launch by US, EU and Partners of the Global Methane Pledge. European Commission - European Commission.

4. Saunois M, Stavert AR, Poulter B et al. The Global Methane Budget 2000–2017. Earth System Science Data 2020;12:1561–623.

5. Tian H, Lu C, Ciais P et al. The terrestrial biosphere as a net source of greenhouse gases to the atmosphere. Nature 2016;531:225–8.

6. Conrad R. Importance of hydrogenotrophic, aceticlastic and methylotrophic methanogenesis for methane production in terrestrial, aquatic and other anoxic environments: A mini review. Pedosphere 2020;30:25–39.

7. Ungerfeld EM. Inhibition of Rumen Methanogenesis and Ruminant Productivity: A Meta- Analysis. Front Vet Sci 2018;5:113.

8. Hook SE, Wright A-DG, McBride BW. Methanogens: Methane Producers of the Rumen and Mitigation Strategies. Archaea 2010;2010:945785.

9. Duin EC, Wagner T, Shima S et al. Mode of action uncovered for the specific reduction of methane emissions from ruminants by the small molecule 3-nitrooxypropanol. Proceedings of the National Academy of Sciences 2016;113:6172–7.

10. Ungerfeld EM. Metabolic Hydrogen Flows in Rumen Fermentation: Principles and Possibilities of Interventions. Front Microbiol 2020;11, DOI: 10.3389/fmicb.2020.00589.

11. Orzuna-Orzuna JF, Dorantes-Iturbide G, Lara-Bueno A et al. Effects of Dietary Tannins’ Supplementation on Growth Performance, Rumen Fermentation, and Enteric Methane Emissions in Beef Cattle: A Meta-Analysis. Sustainability 2021;13:7410.

12. Becker PM, van Wikselaar PG, Franssen MCR et al. Evidence for a hydrogen-sink mechanism of (+)catechin-mediated emission reduction of the ruminant greenhouse gas methane. Metabolomics 2014;10:179–89.

13. Bueno de Mesquita CP, Wu D, Tringe SG. Methyl-Based Methanogenesis: an Ecological and Genomic Review. Microbiology and Molecular Biology Reviews 2023;87:e00024–22.

14. Ellenbogen JB, Borton MA, McGivern BB, et al. Methylotrophy in the Mire: direct and indirect routes for methane production in thawing permafrost. mSystems 2023;9:e00698–23.

15. Khairunisa BH, Heryakusuma C, Ike K et al. Evolving understanding of rumen methanogen ecophysiology. Front Microbiol 2023;14, DOI: 10.3389/fmicb.2023.1296008.

16. Leahy SC, Janssen PH, Attwood GT et al. Electron flow: key to mitigating ruminant methanogenesis. Trends in Microbiology 2022;30:209–12.

17. McGivern BB, Tfaily MM, Borton MA et al. Decrypting bacterial polyphenol metabolism in an anoxic wetland soil. Nature Communications 2021;12:2466.

18. Fofana A, Anderson D, McCalley CK et al. Mapping substrate use across a permafrost thaw gradient. Soil Biology and Biochemistry 2022;175:108809.

19. Parada AE, Needham DM, Fuhrman JA. Every base matters: assessing small subunit rRNA primers for marine microbiomes with mock communities, time series and global field samples. Environmental Microbiology 2016;18:1403–14.

20. Apprill A, McNally S, Parsons R et al. Minor revision to V4 region SSU rRNA 806R gene primer greatly increases detection of SAR11 bacterioplankton. Aquatic Microbial Ecology 2015;75:129–37.

21. Caporaso JG, Lauber CL, Walters WA et al. Ultra-high-throughput microbial community analysis on the Illumina HiSeq and MiSeq platforms. ISME Journal 2012;6:1621–4.

22. Bolyen E, Rideout JR, Dillon MR et al. Reproducible, interactive, scalable and extensible microbiome data science using QIIME 2. Nat Biotechnol 2019;37:852–7.

23. Callahan BJ, McMurdie PJ, Rosen MJ et al. DADA2: High-resolution sample inference from Illumina amplicon data. Nature Methods 2016;13:581–3.

24. Chaumeil PA, Mussig AJ, Hugenholtz P et al. GTDB-Tk: A toolkit to classify genomes with the genome taxonomy database. Bioinformatics 2020;36:1925–7.

25. Oksanen J, Simpson GL, Blanchet FG et al. vegan: Community Ecology Package. 2024.

26. Joshi N, Fass J. Sickle: A sliding-window, adaptive, quality-based trimming tool for FastQ files (Version 1.33) [Software]. Available at https://github.com/najoshi/sickle 2011:2011.

27. Li D, Liu C-M, Luo R et al. MEGAHIT: an ultra-fast single-node solution for large and complex metagenomics assembly via succinct de Bruijn graph. Bioinformatics 2015;31:1674–6.

28. Bushnell B. BBtools. https://jgi.doe.gov/data-and-tools/software-tools/bbtools/.

29. Li H, Handsaker B, Wysoker A et al. The Sequence Alignment/Map format and SAMtools. Bioinformatics 2009;25:2078–9.

30. Kang DD, Li F, Kirton E et al. MetaBAT 2: An adaptive binning algorithm for robust and efficient genome reconstruction from metagenome assemblies. PeerJ 2019;2019:e7359.

31. Chklovski A, Parks DH, Woodcroft BJ et al. CheckM2: a rapid, scalable and accurate tool for assessing microbial genome quality using machine learning. Nat Methods 2023;20:1203–12.

32. Bowers RM, Kyrpides NC, Stepanauskas R et al. Minimum information about a single amplified genome (MISAG) and a metagenome-assembled genome (MIMAG) of bacteria and archaea. Nature Biotechnology 2017;35:725–31.

33. Olm MR, Brown CT, Brooks B et al. DRep: A tool for fast and accurate genomic comparisons that enables improved genome recovery from metagenomes through de-replication. ISME Journal 2017;11:2864–8.

34. McGivern BB, Cronin DR, Ellenbogen JB et al. Microbial polyphenol metabolism is part of the thawing permafrost carbon cycle. Nat Microbiol 2024;9:1454–66.

35. Chaumeil P-A, Mussig AJ, Hugenholtz P et al. GTDB-Tk v2: memory friendly classification with the genome taxonomy database. Bioinformatics 2022;38:5315–6.

36. Shaffer M, Borton MA, McGivern BB et al. DRAM for distilling microbial metabolism to automate the curation of microbiome function. Nucleic acids research 2020;48:8883–900.

37. Langmead B, Salzberg SL. Fast gapped-read alignment with Bowtie 2. Nature Methods 2012;9:357–9.

38. Anders S, Pyl PT, Huber W. HTSeq--a Python framework to work with high-throughput sequencing data. Bioinformatics 2015;31:166–9.

39. Smid M, Coebergh van den Braak RRJ, van de Werken HJG et al. Gene length corrected trimmed mean of M-values (GeTMM) processing of RNA-seq data performs similarly in intersample analyses while improving intrasample comparisons. BMC Bioinformatics 2018;19:236.

40. Anantharaman K, Brown CT, Hug LA et al. Thousands of microbial genomes shed light on interconnected biogeochemical processes in an aquifer system. Nat Commun 2016;7:13219.

41. Søndergaard D, Pedersen CNS, Greening C. HydDB: A web tool for hydrogenase classification and analysis. Sci Rep 2016;6:34212.

42. Portman TA, Arnold AE, Bradley RG et al. Fungal endophytes of the invasive grass *Eragrostis lehmanniana* shift metabolic expression in response to native and invasive grasses. Fungal Ecology 2024;68:101327.

43. Wyatt M, Choudhury A, Von Dohlen G et al. Randomized control trial of moderate dose vitamin D alters microbiota stability and metabolite networks in healthy adults. Microbiology Spectrum 2024;0:e00083–24.

44. Sumner LW, Amberg A, Barrett D et al. Proposed minimum reporting standards for chemical analysis Chemical Analysis Working Group (CAWG) Metabolomics Standards Initiative (MSI). Metabolomics 2007;3:211–21.

45. R: The R Project for Statistical Computing.

46. Wickham H. Ggplot2. 2016th ed. Springer-Verlag New York

47. Wickham H, Vaughan D, Girlich M, et al. tidyr: Tidy Messy Data. 2023.

48. Wickham H, François R, Henry L, et al. dplyr: A Grammar of Data Manipulation. 2022.

49. Kolde R. pheatmap: Pretty Heatmaps. 2019.

50. Matsuda M, Otsuka Y, Jin S et al. Biotransformation of (+)-catechin into taxifolin by a two- step oxidation: Primary stage of (+)-catechin metabolism by a novel (+)-catechin-degrading bacteria, Burkholderia sp. KTC-1, isolated from tropical peat. Biochemical and Biophysical Research Communications 2008;366:414–9.

51. Otsuka Y, Matsuda M, Sonoki T et al. Enzymatic activity of cell-free extracts from Burkholderia oxyphila OX-01 bio-converts (+)-catechin and (−)-epicatechin to (+)-taxifolin. Bioscience, Biotechnology, and Biochemistry 2016;80:2473–9.

52. Braune A, Gütschow M, Blauta M. An NADH-dependent reductase from Eubacterium ramulus catalyzes the stereospecific heteroring cleavage of flavanones and flavanonols. Applied and Environmental Microbiology 2019;85:1233–52.

53. Braune A, Engst W, Elsinghorst PW et al. Chalcone isomerase from Eubacterium ramulus catalyzes the ring contraction of flavanonols. Journal of Bacteriology 2016;198:2965–74.

54. Braune A, Gütschow M, Engst W et al. Degradation of Quercetin and Luteolin byEubacterium ramulus. Applied and Environmental Microbiology 2001;67:5558–67.

55. Maini Rekdal V, Nol Bernadino P, Luescher MU et al. A widely distributed metalloenzyme class enables gut microbial metabolism of host- and diet-derived catechols. Elife 2020;9:e50845.

56. Zhou Y, Wei Y, Jiang L et al. Anaerobic phloroglucinol degradation by Clostridium scatologenes. mBio 2023;14:e01099–23.

57. Brune A, Schink B. Pyrogallol-to-phloroglucinol conversion and other hydroxyl-transfer reactions catalyzed by cell extracts of Pelobacter acidigallici. Journal of Bacteriology 1990;172:1070–6.

58. Mondav R, Woodcroft BJ, Kim E-H et al. Discovery of a novel methanogen prevalent in thawing permafrost. Nat Commun 2014;5:3212.

59. Woodcroft BJ, Singleton CM, Boyd JA et al. Genome-centric view of carbon processing in thawing permafrost. Nature 2018;560:49–54.

60. Wintsche B, Jehmlich N, Popp D et al. Metabolic Adaptation of Methanogens in Anaerobic Digesters Upon Trace Element Limitation. Front Microbiol 2018;9, DOI: 10.3389/fmicb.2018.00405.

61. Markovic O, Janecek S. Pectin methylesterases: sequence-structural features and phylogenetic relationships. Carbohydr Res 2004;339:2281–95.

62. Kotani T, Yurimoto H, Kato N et al. Novel Acetone Metabolism in a Propane-Utilizing Bacterium, Gordonia sp. Strain TY-5. Journal of Bacteriology 2007;189:886–93.

63. Kurth JM, Op den Camp HJM, Welte CU. Several ways one goal—methanogenesis from unconventional substrates. Appl Microbiol Biotechnol 2020;104:6839–54.

64. Krivushin KV, Shcherbakova VA, Petrovskaya LE et al. Methanobacterium veterum sp. nov., from ancient Siberian permafrost. International Journal of Systematic and Evolutionary Microbiology 2010;60:455–9.

65. Borrel G, Joblin K, Guedon A et al. Methanobacterium lacus sp. nov., isolated from the profundal sediment of a freshwater meromictic lake. International Journal of Systematic and Evolutionary Microbiology 2012;62:1625–9.

66. Vignais PM, Billoud B. Occurrence, Classification, and Biological Function of Hydrogenases: An Overview. Chem Rev 2007;107:4206–72.

67. Westerholm M, Calusinska M, Dolfing J. Syntrophic propionate-oxidizing bacteria in methanogenic systems. FEMS Microbiology Reviews 2022;46:fuab057.

