## Supplementary Online Material for "Polyphenol rewiring of the microbiome reduces methane emissions"

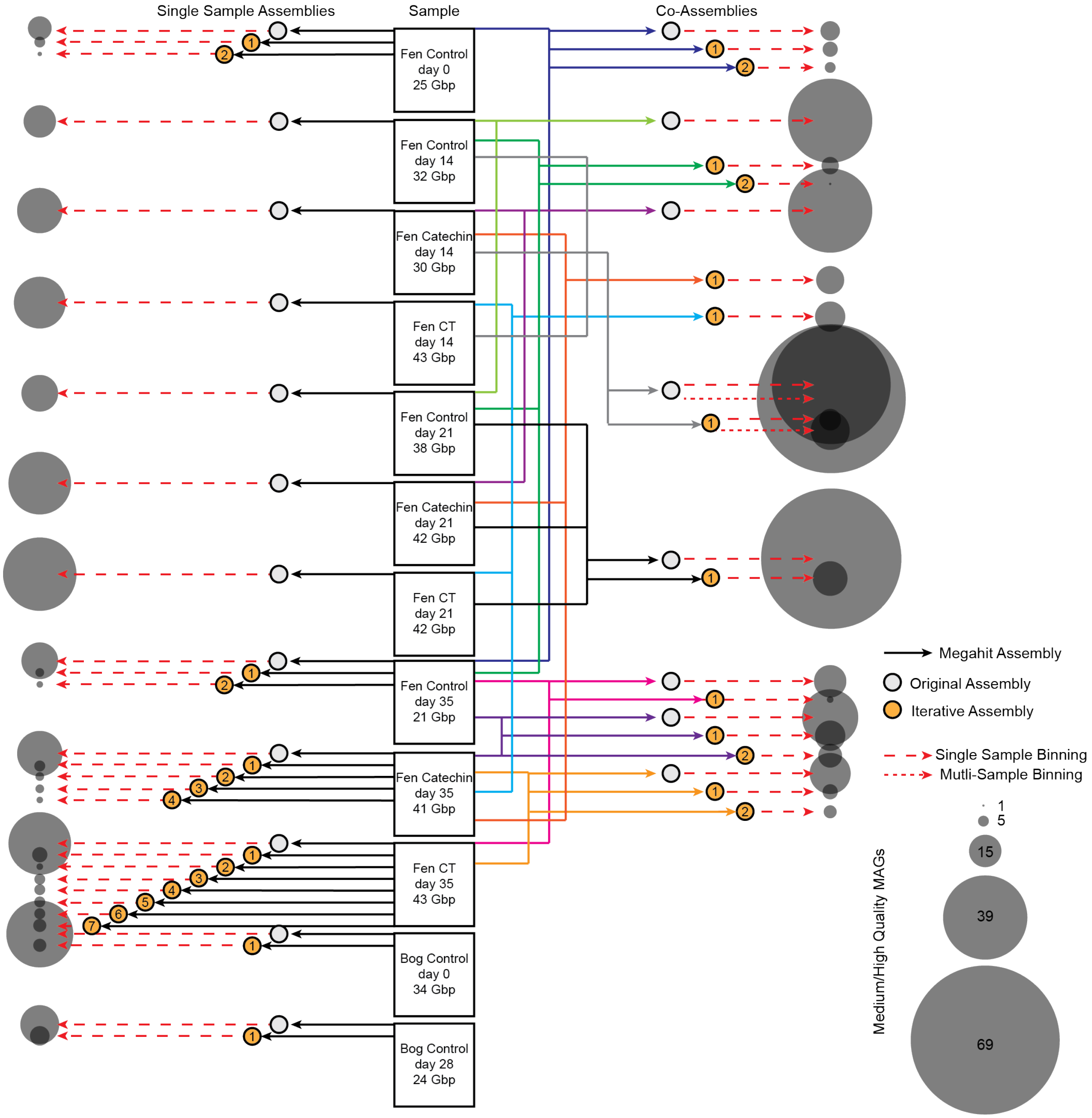


**Supplementary Figure 1.** Workflow for metagenome-assembled genome (MAG) recovery. Twelve metagenomes were obtained (listed in the center) from Stordalen Mire microcosms. Metagenomic reads from these samples were assembled in four ways: (1) single sample assemblies (left, grey circles), (2) single sample iterative assemblies (left, yellow circles), (3) co-assemblies (right, grey circle), (4) iterative co-assemblies (right, yellow circles). Assemblies were binned in two ways: (1) using coverage from a single sample (bigger dashed lines) and (2) using coverage across multiple samples (smaller dashed lines). The number of MAGs recovered from each assembly and binning strategy are represented according to the size of the circles, ranging from 1 (smallest) to 69 (biggest). For metagenome and assembly information, see **Supplementary Data 2.**

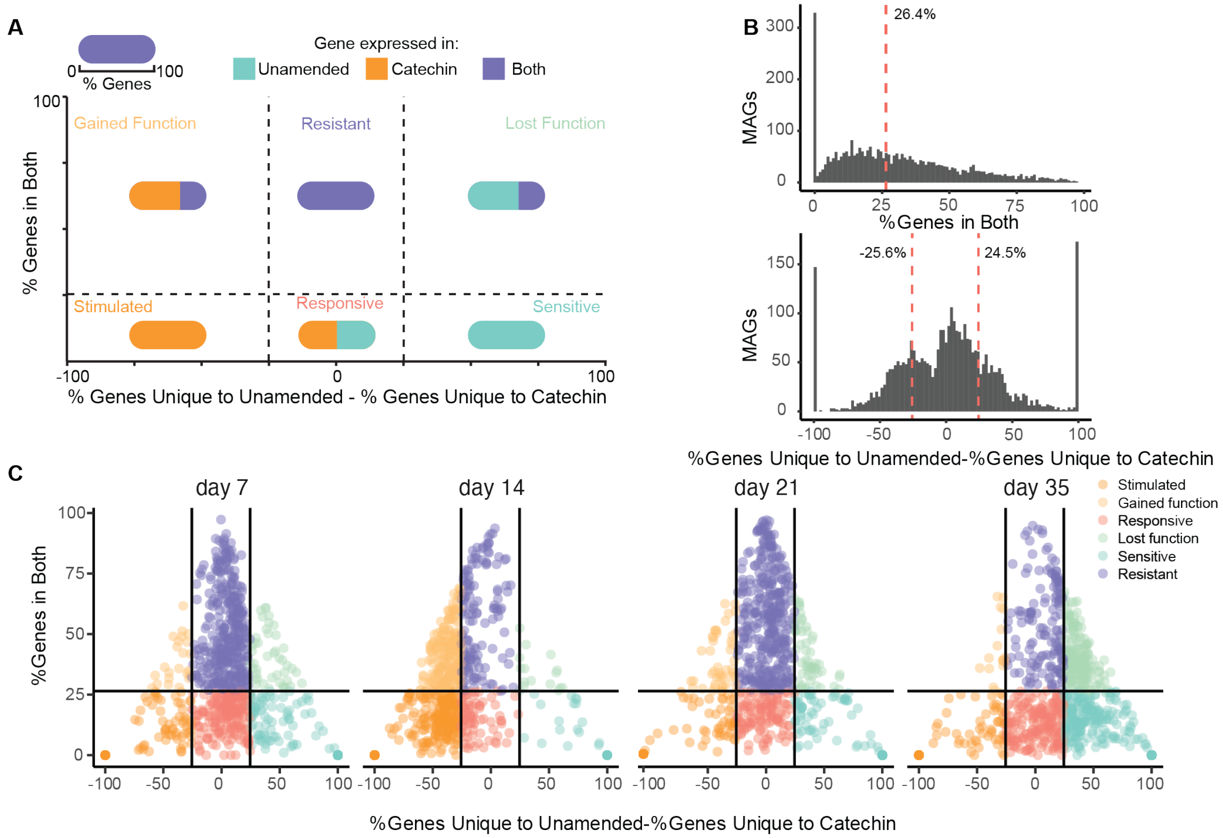


**Supplementary Figure 2. (A)** Schematic of the classification scheme used to describe MAG responses to catechin amendment. **(B)** Thresholds used to assign MAGs to response categories were determined from median value of the percent genes expressed in both treatments (top, used as y-axis threshold in **A**) and the 25% and 75% quantile values of the difference in genes unique to each treatment (bottom, used as x-axis thresholds in **A**). The values for these thresholds are printed on the plots. (**C**) Plots of MAG classification at each timepoint. Each point is a MAG, colored by its classification.

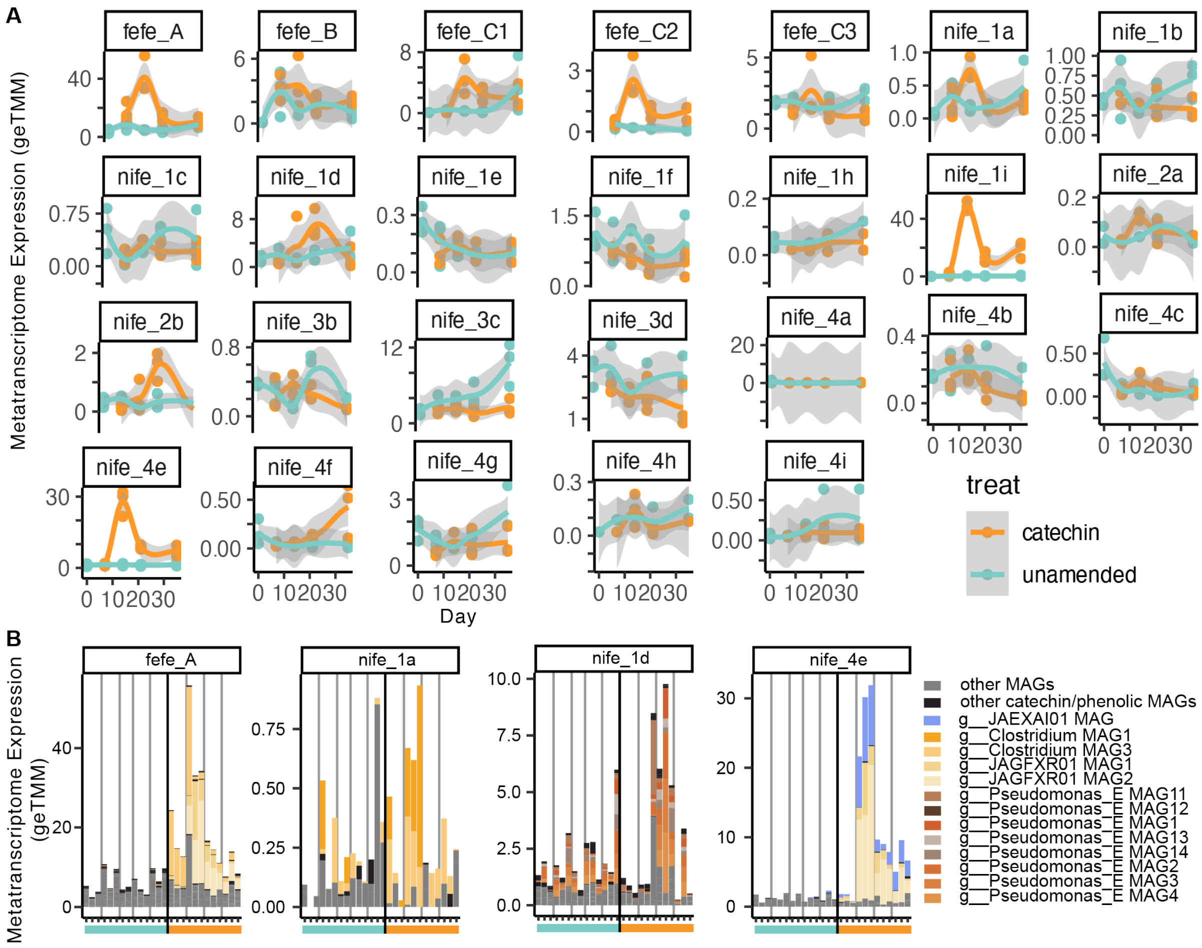


**Supplementary Figure 3. Hydrogenase gene expression.** (A) Summed metatranscriptome expression of hydrogenase genes by hydrogenase subgroup. Smoothed curves represent the average metatranscriptome expression and concentration, respectively(n=3), with individual replicates plotted as points, and the shaded area represents the 95% confidence interval. (B) Stacked barplots of hydrogenase expression by MAG, where coloring corresponds to MAG genus.

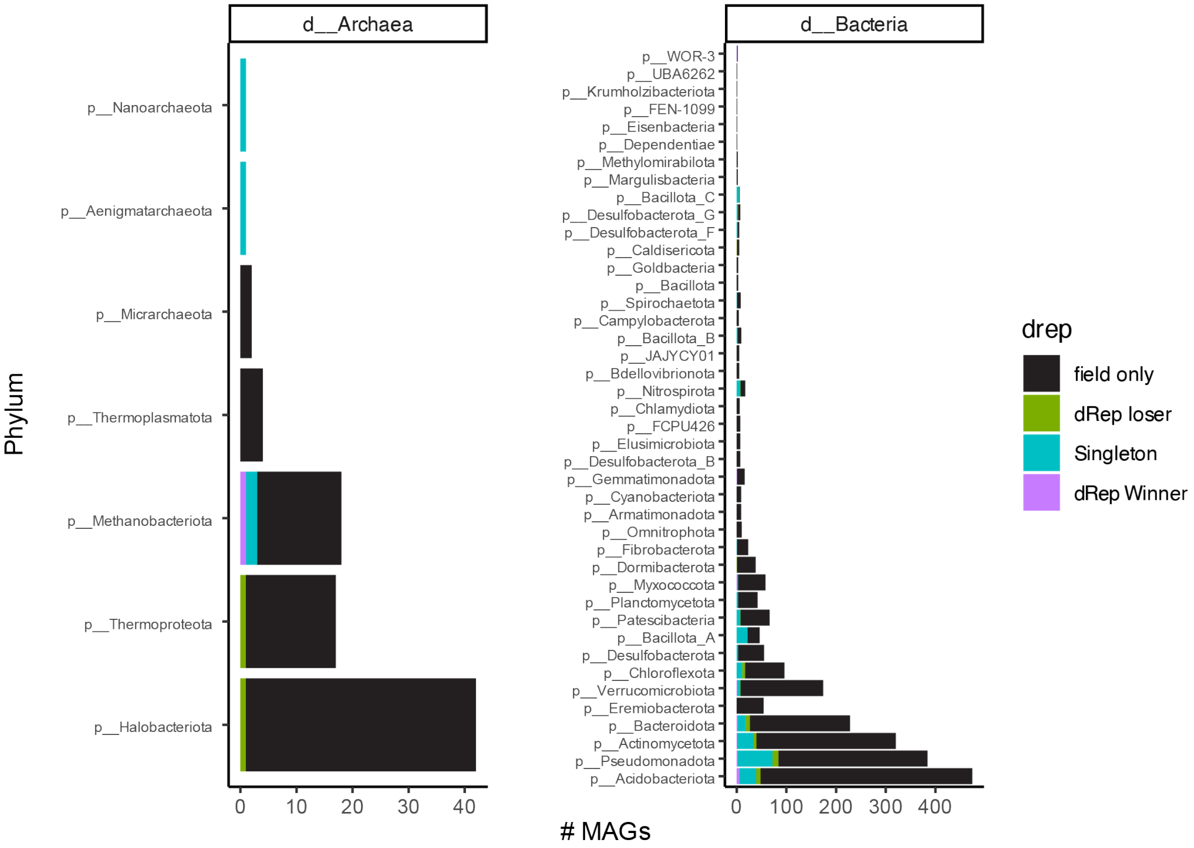


**Supplementary Figure 4.** Phylum level summary of metagenome-assembled genomes (MAGs) in the combined microcosm and field database. The final database was 2,303 MAGs after dereplicating the field MAGs with microcosm MAGs. Bars are colored by whether dRep winners were from clusters of only field MAGs (black), represented by a microcosm MAG (purple, dRep Winner), represented by a field MAG with a microcosm MAG in the cluster (green, dRep loser), or singleton microcosm MAGs (blue, Singleton). For detailed MAG information, see **Supplementary Data 2**.

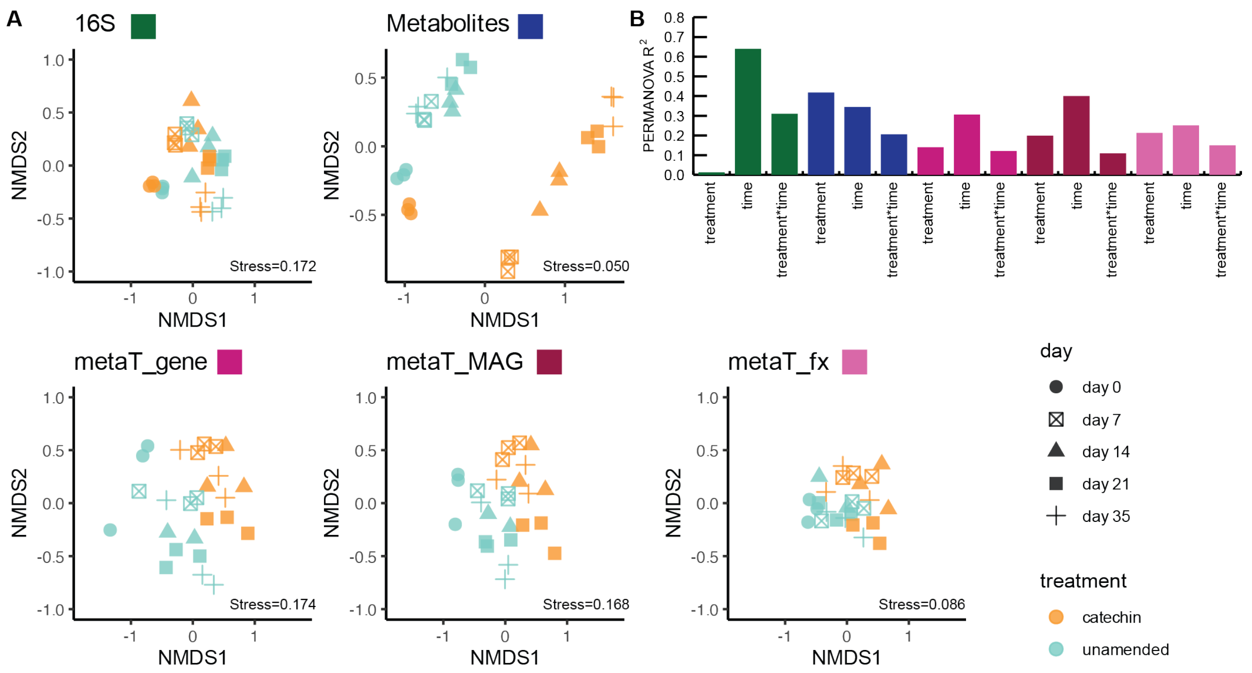


**Supplementary Figure 5. (A)** Non-metric multidimensional scaling (NMDS) ordinations of microbial community measurements. NMDS ordinations of Bray-Curtis distances for 16S rRNA gene amplicon (16S), LC-MS (metabolites), and metatranscriptome data at the gene level (metaT_gene), the genome level (metaT_MAG), and function level (metaT_fx). Points are shaped by timepoint according to the legend at right and colored by treatment. The stress of the ordination is given in the bottom right corner of each ordination. **(B)** The PERMANOVA $R^2$ values of the tested variables time, treatment, and the interaction variable (treatment*time). All tests were significant ($p<0.05$).

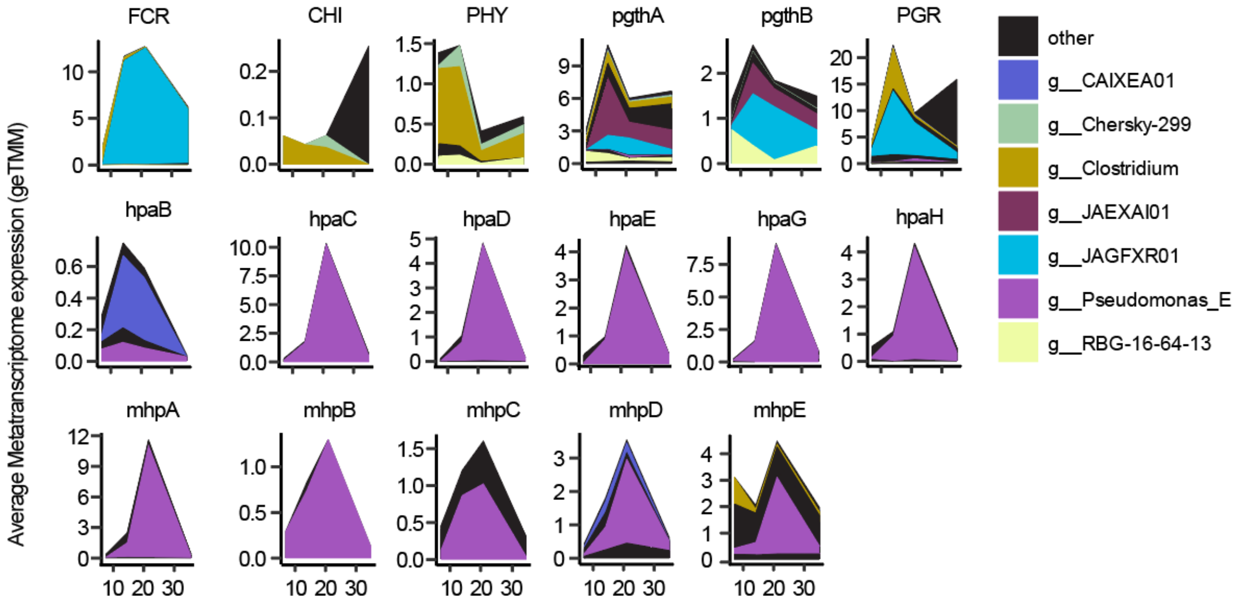


**Supplementary Fig. 6.** Average metatranscriptome expression (n=3) of catechin, phloroglucinol, and phenolic acid degradation genes colored by expressing genus.

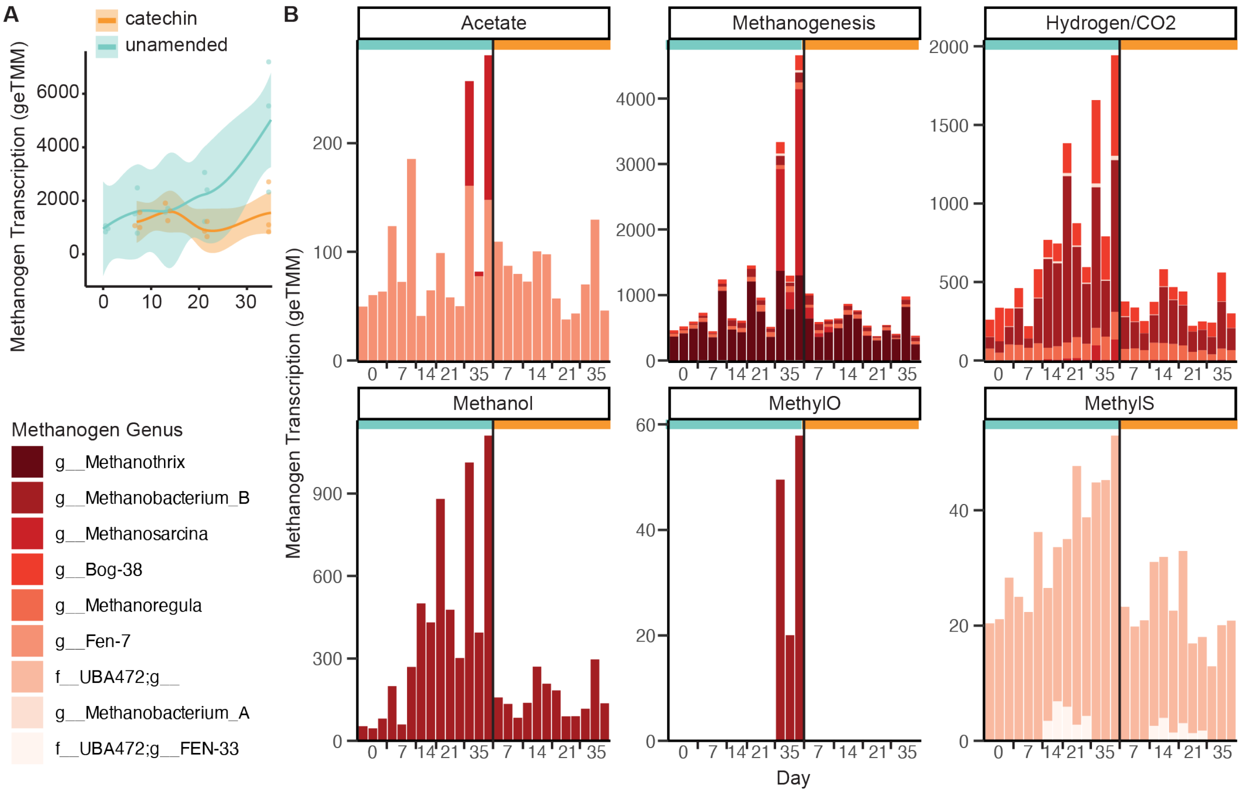


**Supplementary Figure 7. (A)** Summed metatranscriptome expression of methanogen MAGs. Smoothed curves represent the average metatranscriptome expression and concentration, respectively(n=3), with individual replicates plotted as points, and the shaded area represents the 95% confidence interval. **(B)** Relative expression of methanogen MAGs by substrate use over time. MAGs expressing methanogenesis pathway genes, but not substrate specific genes, were classified as “methanogenesis”. Each replicate is plotted as a stacked bar plot, colored by MAG genus. Samples are listed by treatment in chronological order, starting with unamended (teal, left of black line) then catechin-amended (orange, right of black line).

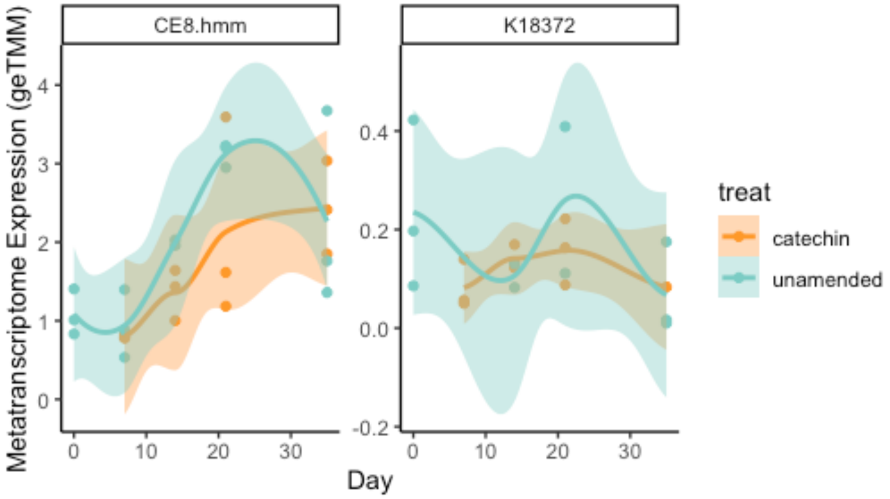


**Supplementary Figure 8.** Summed metatranscriptome expression of pectin methylesterase (CE8.hmm, methanol from pectin) and methyl acetate acetohydrolase (K18372, methanol from acetone). Smoothed curves represent the average metatranscriptome expression and concentration, respectively (n=3), with individual replicates plotted as points, and the shaded area represents the 95% confidence interval.

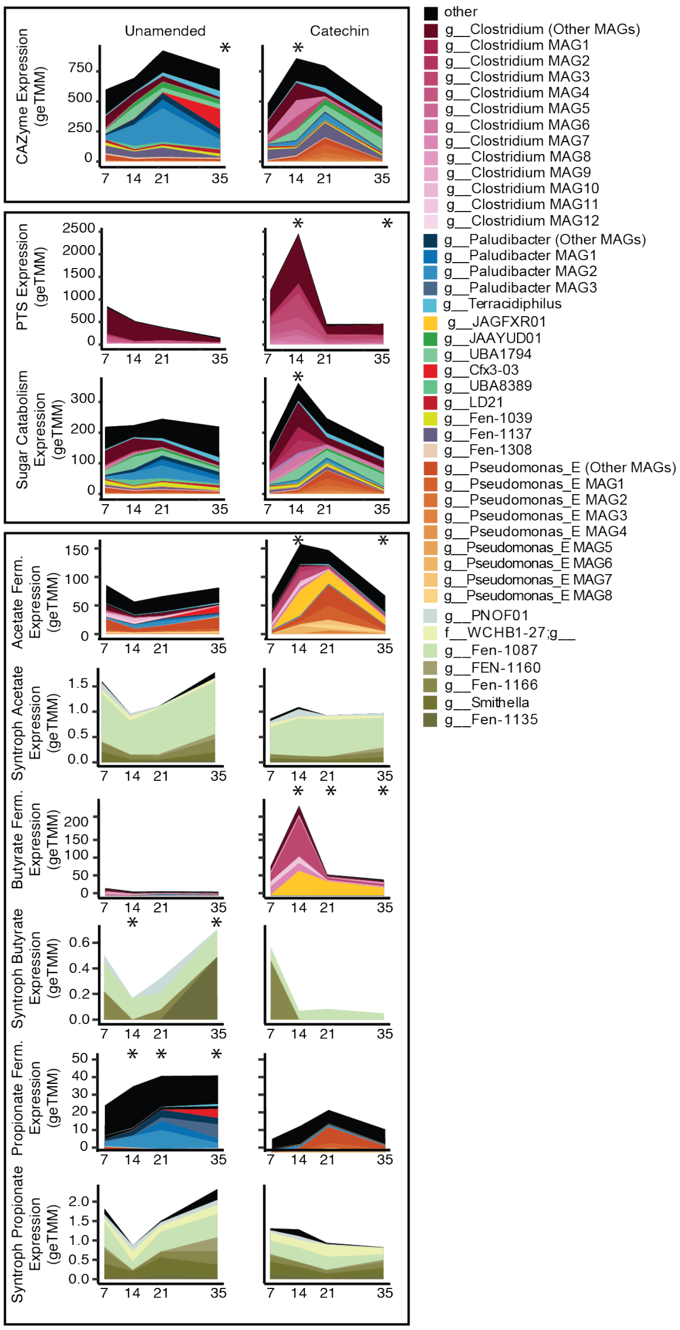


**Supplementary Figure 9.** Area plots showing the average gene expression (n=3) at each time point for specified pathways overtime, with colored slices corresponding to the expressing lineages.

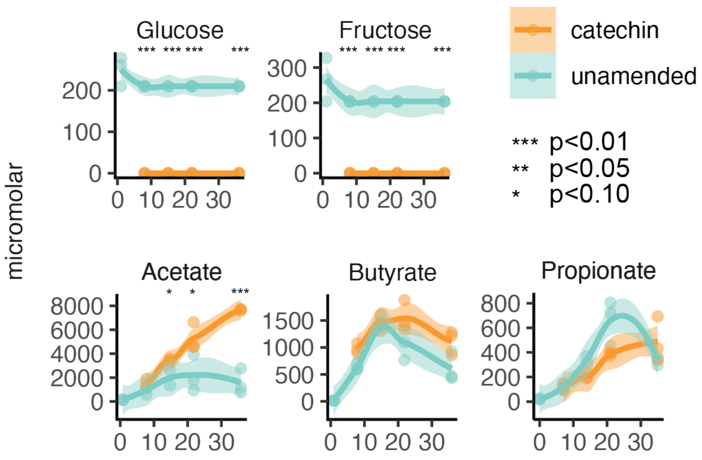


**Supplementary Figure 10.** Plots of the concentration (micromolar) of various metabolites detected by NMR. Smoothed curves represent the average metatranscriptome expression and concentration, respectively (n=3), with individual replicates plotted as points, and the shaded area represents the 95% confidence interval. Timepoints with significant differences between unamended and catechin-amended are marked with asterisks (two-sided t-test, * p-value <0.10, ** p-value <0.05, *** p-value <0.01).

**Supplementary Table 1.** Catechin response categories.

| **Response to catechin** | **% Genes Expressed in Both Conditions (%both)** | **Difference in % of treatment-unique genes (Δunique)** | **Example** |
|---|---|---|---|
| Resistant | >26% of genes expressed in both treatments | \|Δunique\| <25% difference in the gene expression | A MAG expressed 100 total genes at day 7. Of these 100 genes, 30 were expressed in both unamended and catechin treatments (%both=30%), 40 were only in unamended, and 30 were only in catechin (Δunique = 40-30=10%). |
| Responsive | <26% of genes expressed in both treatments | \|Δunique\| <25% difference in the gene expression | A MAG expressed 100 total genes at day 7. Of these 100 genes, 20 were expressed in both unamended and catechin treatments (%both=20%), 50 were only in unamended, and 30 were only in catechin (Δunique = 50-30=20%). |
| Sensitive | <26% of genes expressed in both treatments | Δunique >25% difference in the gene expression | A MAG expressed 100 total genes at day 7. Of these 100 genes, 20 were expressed in both unamended and catechin treatments (%both=20%), 70 were only in unamended, and 10 were only in catechin (Δunique = 70-10=60%). |
| Stimulated | <26% of genes expressed in both treatments | Δunique <-25% difference in the gene expression | A MAG expressed 100 total genes at day 7. Of these 100 genes, 20 were expressed in both unamended and catechin treatments (%both=20%), 10 were only in unamended, and 70 were only in catechin (Δunique = 10-70=-60%). |
| Lost Function | >26% of genes expressed in both treatments | Δunique >25% difference in the gene expression | A MAG expressed 100 total genes at day 7. Of these 100 genes, 30 were expressed in both unamended and catechin treatments (%both=30%), 60 were only in unamended, and 10 were only in catechin (Δunique = 60-10=-50%). |
| Gained Function | >26% of genes expressed in both treatments | Δunique<-25% difference in the gene expression | A MAG expressed 100 total genes at day 7. Of these 100 genes, 30 were expressed in both unamended and catechin treatments (%both=30%), 10 were only in unamended, and 60 were only in catechin (Δunique = 10-60=-50%). |

**Supplementary Table 2.**

| Syntrophic Pathway | Reaction | $\Delta G^{\circ\prime}$ kJ/mol | $\Delta G$ | Hypothesized Change under Catechin Amendment |
|---|---|---|---|---|
| **Propionate Oxidation via mmc** | Propionate + $2H_2O$ → Acetate + $CO_2$ + $3H_2$ | +73.7 | $\Delta G^{\circ\prime} + RT\ln(\frac{[acetate][CO_2][H_2]^3}{[propionate]})$ | Less favorable<br>• Acetate ↑ (NMR) = decrease favorability<br>• $H_2$ ↓ (hydrogenases) = increase favorability<br>• Propionate unchanged (NMR) = no impact<br>• No data on dissolved $CO_2$ |
| **Propionate Oxidation via dismutation** | 2Propionate → Acetate + Butyrate | +2.8 | $\Delta G^{\circ\prime} + RT\ln(\frac{[acetate][butyrate]}{[propionate]})$ | Less favorable<br>• Acetate ↑ (NMR) = decrease favorability<br>• Butyrate unchanged (NMR) = no impact<br>• Propionate unchanged (NMR) = no impact |

| | | | | |
|---|---|---|---|---|
| **Acetate Oxidation** | Acetate + $H^+$ + $2H_2O$ → $2CO_2$ + $4H_2$ | +94.9 | $\Delta G^{\circ\prime} + RT\ln(\frac{[CO_2]^2[H_2]^4}{[acetate][H^+]})$ | More favorable<br>• No data on dissolved $CO_2$<br>• $H_2$ ↓ (hydrogenases) = increase favorability<br>• Acetate ↑ (NMR) = increase favorability<br>• $H^+$ ↑ (pH) = increase favorability |
| **Butyrate Oxidation** | Butyrate + $2H_2O$ → 2Acetate + $2H_2$ + $H^+$ | +49.8 | $\Delta G^{\circ\prime} + RT\ln(\frac{[acetate][H_2]^2[H^+]}{[butyrate]})$ | Less favorable<br>• Acetate ↑ (NMR) = decrease favorability<br>• $H_2$ ↓ (inferred from hydrogenases) = increase favorability<br>• $H^+$ ↑ (pH) = decrease favorability<br>• Butyrate unchanged (NMR) = no impact |
